# A computational analysis of the glycoprotein LRP1 structure and the role of glycans as quaternary glue

**DOI:** 10.1101/2025.06.10.658798

**Authors:** Gian Marco Tuveri, Marco Basile, Silvia Acosta Gutiérrez, Marius Kausas, Silvia Pujals, Xiaohe Tian, Giancarlo Franzese, Lorena Ruiz Pérez, Giuseppe Battaglia

**Affiliations:** Institute for Bioengineering of Catalunya (IBEC), The Barcelona Institute of Science and Technology (BIST), Barcelona, Spain; Institute of Nanoscience and Nanotechnology (IN2UB), University of Barcelona, Barcelona, Spain; Department of Condensed Matter Physics, University of Barcelona, Barcelona, Spain; Department of Biomedicine, University of Barcelona, Barcelona, Spain; Vianautis Bio ltd, Cambridge, United Kingdom; Department of Biological Chemistry, Institute for Advanced Chemistry of Catalonia, Barcelona, Spain; Department of Radiology and Huaxi MR Research Center (HMRRC), Functional and Molecular Imaging Key Laboratory of Sichuan Province, Department of Radiology and National Clinical Research Center for Geriatrics, Frontiers Science Center for Disease-Related Molecular Network, State Key Laboratory of Biotherapy, West China Hospital of Sichuan University, Chengdu, 610000, Sichuan Province, China; Serra Húnter Fellow, Department of Applied Physics, University of Barcelona, Barcelona, Spain; Catalan Institution for Research and Advanced Studies (ICREA), Barcelona, Spain

## Abstract

The low-density lipoprotein receptor-related protein 1 (LRP1) plays a critical role in development and transport across the blood-brain barrier (BBB), yet until now, its molecular architecture remained unresolved due to the absence of an experimentally determined structure. Using homology modeling and neural network-based structure prediction algorithms, complemented with molecular dynamics (MD) simulations, we propose a comprehensive model of LRP1 structures. We observe a natural dimerization mechanism and provide insight into the dynamic behavior of its flexible domains under physiological conditions. We investigated the stability of non-covalent interactions keeping LRP1’s α and β chains linked together, and found the energy required to break the link is 180±2 kT. The MD characterization highlights the fundamental role of glycans in the creation of LRP1’s quaternary structure, increasing the number of intra-dimeric contacts. This study opens new avenues for targeted drug design strategies, enhancing our molecular understanding of LRP1’s receptor-mediated transport in the brain and the key mediation of glycosylation in protein-protein interactions.

## Introduction

The low-density lipoprotein (LDL) receptor-related protein [1, 2] (LRP1) is a multifunctional glycoprotein involved in diverse biological pathways, including cancer, infection, and neurodegeneration. LRP1 is expressed in many animal tissues, and its malfunction affects various pathways in humans, causing cancer, autoimmune, neurodegenerative, and cardiovascular diseases [1–6]. LRP1 is a critical gene, and its deletion prevents vascular and neural development [7, 8]. As a key endocytic and signaling receptor, LRP1 regulates cellular homeostasis, metabolism, and immune responses, making it a critical player in health and disease [1, 2, 9]. Notably, LRP1 can interact with over forty different ligands [1, 2, 9]. This broad ligand-binding capacity may be attributed to its considerable size, comprising 4,544 amino acids, which allows for multiple interaction domains. LRP1 is initially synthetized as a single chain precursor and further processed by furin in the trans-Golgi compartment, resulting in a non-covalently associated heterodimer composed of a 515 kDa extracellular α-chain and an 85 kDa transmembrane β-chain [1, 9, 10]. The extracellular portion of LRP1, or ectodomain, was observed in 2017 by De Nardis and collaborators [11] with negative-stain electron microscopy and small angle x-ray scattering (SAXS), providing the first structural insights into its overall shape and flexibility in solution. All the images showed a maximum extension of 35nm and a monomeric structure. More accurate observations of LRP1 and other members of the LDL receptor family reported the structure of the three motifs or components of the protein (see Fig. 1). The first motif is the calcium binding (CB) motif, believed to constitute the binding domains of LRP1 and to have two main features: the conservation of six cysteines creating three sulfur bridges and four to six acidic amino acids creating an electrostatic favorable coordination site for a coordinated calcium ion [12, 13]. The second is the β-propeller motif [14], formed of six β sheets interacting by hydrophobic interactions and with the characteristic propeller shape. Finally, the epithelial growth factor (EGF)-like motif [15], which is also stabilized by three sulfur bridges does not show a stable coordination site for an ion.

**Figure 1.**
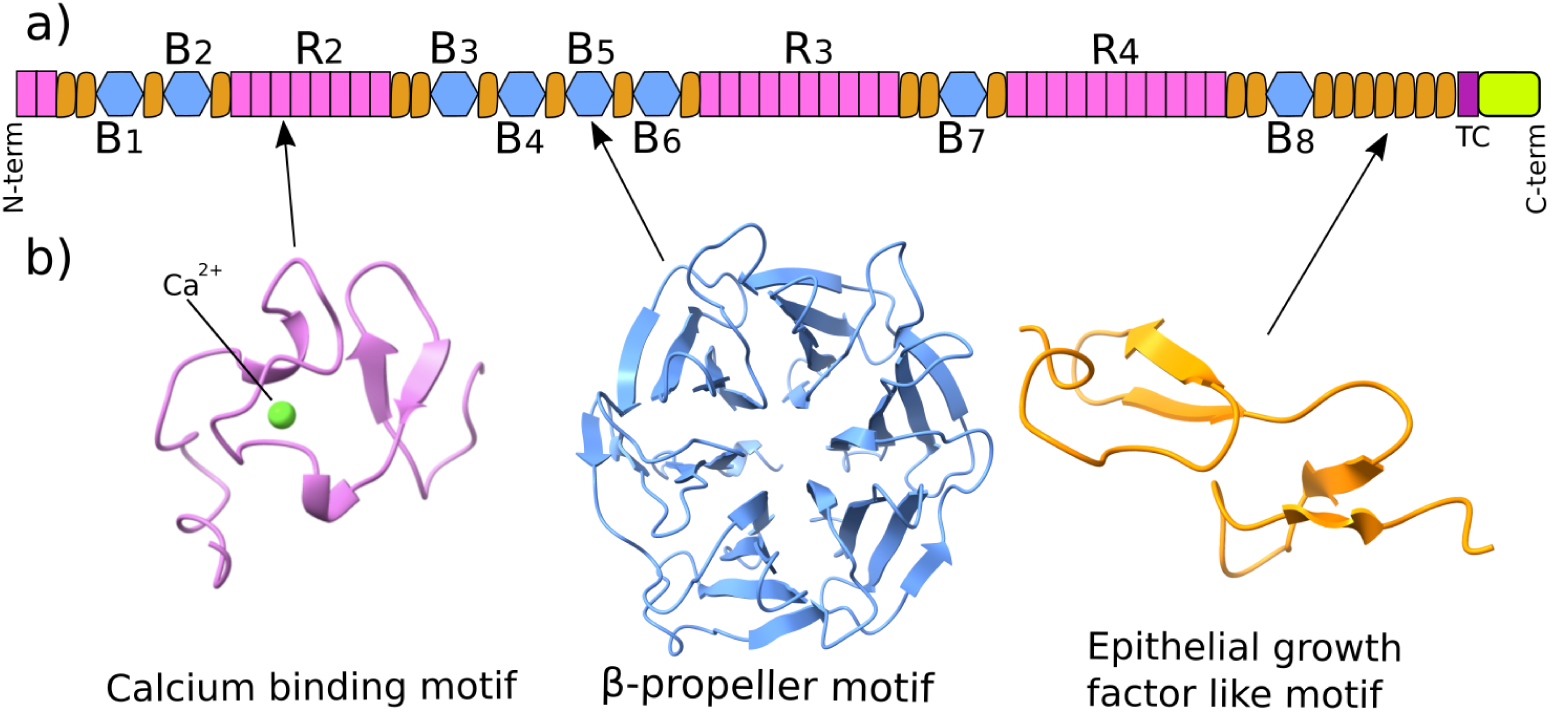
a) The colored scheme shows the motif sequence in LRP1. Each color corresponds to a different structural motif: the calcium-binding (pink), β-propeller (blue), and EGF-like (orange) domains found in the LRP family. b) The three structural motifs observed in another member of the LDL receptor family: LRP2 (8EM4).

While the primary sequence has been known since the discovery of the receptor [16], together with its local secondary and local tertiary structures, the scientific community has not produced a satisfyingly accurate characterization of full LRP1’s tertiary structure. To understand the latter better, it is useful to refer to the structures of homologous proteins. Recently, the structure of LRP2, a member of the LDL receptor family [17], was resolved using cryo-electron microscopy (PDB ID: 8EM4) [18]. LRP2, also known as megalin, shares the ability to bind numerous ligands with LRP1, some of which are common to both receptors [19]. Comprising 4655 amino acids, LRP2 features the same structural motifs as other LDL receptor family members, including LRP1, with conserved motif order over long stretches of the protein chains [19]. The resolved LRP2 structure reveals it as a homodimer, with two monomers interacting through hydrogen bonds and salt bridges in specific regions, leading to a quaternary structure; a following work confirmed the structure at physiological pH [20]. This homodimer configuration offers exciting new insights into molecular recognition, binding mechanisms, and transport processes. The similarity between LRP2 and LRP1 raises the possibility that LRP1 might also function in a dimeric state.

Both LRP1 and LRP2 are glycoproteins that are characterized by the presence of covalently attached glycans (complex carbohydrate chains). These glycans play a crucial role in modulating protein-protein interactions, often by stabilizing particular protein conformations or by shielding proteolytic cleavage sites to maintain protein integrity [21]. Increasing evidence has also pointed to the importance of glycan-mediated mechanisms in viral infectivity, most notably in the case of SARS-CoV-2, where glycans influence viral entry and immune evasion [22, 23]. As a result, the field of glycoscience has gained considerable momentum, emerging as a vital area of research with broad implications across molecular biology and biomedical sciences [24].

LRP1 is involved in the transport and clearance of several extracellular proteins, including amyloid beta and tau, both of which are key pathological factors in neurodegenerative diseases such as Alzheimer’s disease. It therefore plays a critical role in maintaining vascular integrity and mediating the removal of toxic species from the brain across the blood–brain barrier (BBB) [25]. In a previous work, we showed that this process involves the creation of tubular vesicular shuttles stabilized by the F-BAR domain protein syndapin-2 [26]. The created tubules allow a fast and effective transcytosis passage of misfolded proteins and synthetic nanoparticles across the BBB [10, 26]. A central concept in our previous work [26] on LRP1 binding is multivalency, which refers to the ability of multiple LRP1 receptors to simultaneously engage with the ligands displayed on a single nanoparticle. The overall binding strength, or avidity, of these multivalent interactions plays a critical role in determining the route of nanoparticle transport across the BBB. As such, avidity is a key parameter to consider when designing new nanoparticles. To optimize this interaction, it is essential to understand how LRP1 interacts with potential ligands that will decorate the nanoparticle surface. This investigation begins with the production of LRP1’s structure.

To further investigate the role of LRP1 in the BBB, in this work, we investigated the localization of LRP1 on the surface of brain endothelial cells and present two novel predictions for the quaternary structures of LRP1: monomeric and dimeric, with the dimeric form derived through homology modeling based on LRP2. These structures were further analyzed for their dynamic properties, emphasizing the role of domain flexibility in the protein, the contribution of glycans to stabilizing the dimeric conformation, and the stability of the non-covalent interactions between the two LRP1 chains.

## Results

### LRP1 co-localizes on the surface of bEnd3 cells

Using stochastic optical reconstruction microscopy (STORM) imaging, we investigated the structural organization of LRP1 on brain endothelial cells with spatial resolution in the tens of nanometers (Fig. 2.a). In our experiments, cells were labelled using an Alexa-647 conjugated secondary antibody, and we first calibrated our system by imaging these antibodies under highly diluted conditions as detailed in the methods section. This calibration, aided by Mean Shift Clustering (MSC) analysis, established that a single secondary antibody produced a median of six fluorescence localizations, setting a baseline to distinguish individual molecules from clusters (Fig. 2.c). When we imaged LRP1 on bEnd3 cell membranes via immunofluorescence, we observed that, unlike the uniformly distributed single antibody signals, LRP1 was organized into discrete clusters, with some clusters exhibiting up to 30 fluorescence localizations (Fig. 2.b). Further validation through varying the clustering bandwidth from 30 to 200 nm confirmed that even larger bandwidths occasionally merged signals, the LRP1 clusters maintained a more complex distribution than the calibrated secondary antibody (Fig. 2.d). This comprehensive analysis confirms that LRP1 forms higher order structures, with multiple molecules colocalizing in small, discrete clusters on the cell surface. This analysis confirms the ability of LRP1 to interact, forming dimer, trimer, and possibly higher order arrangements.

**Figure 2.**
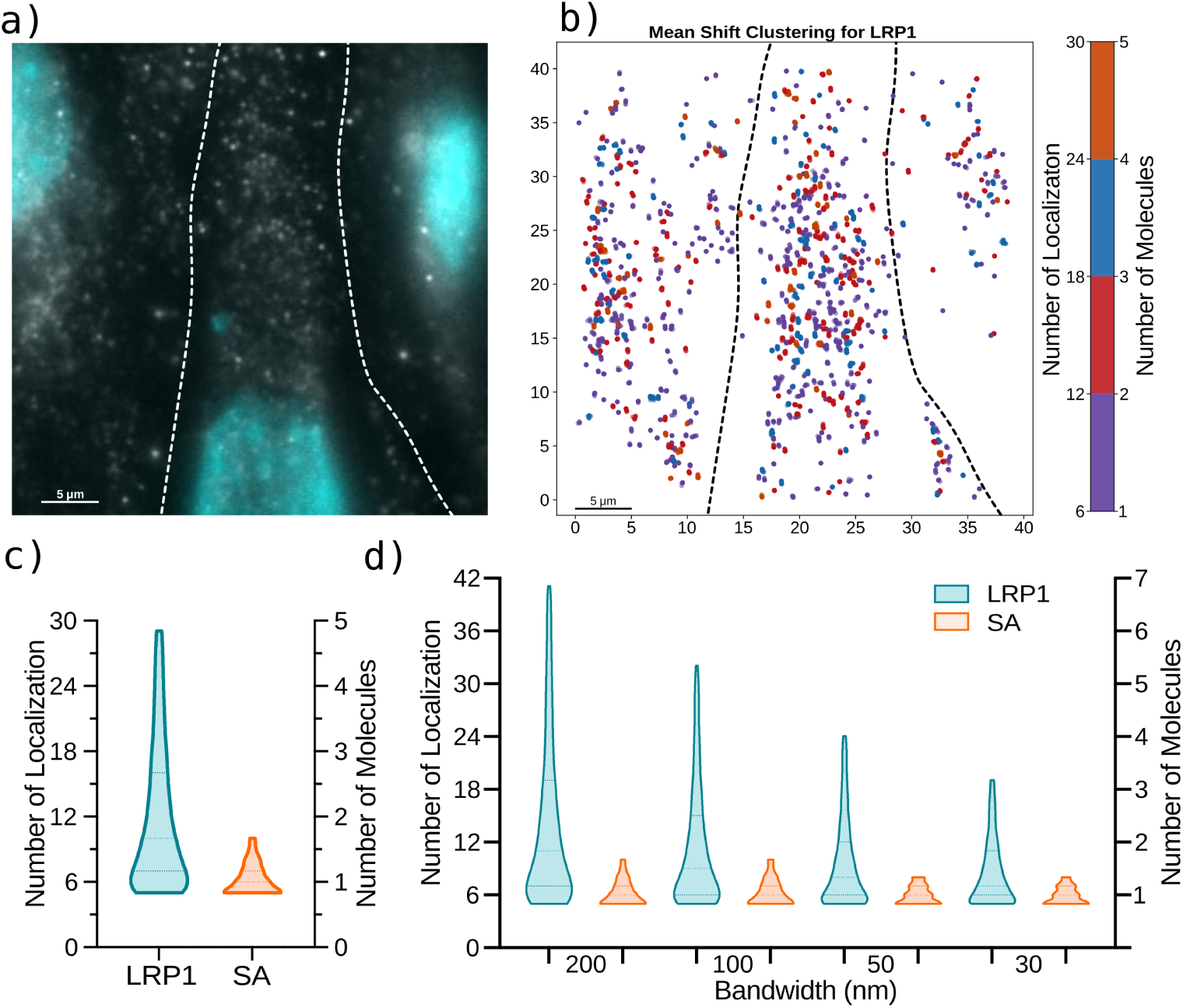
a) Low-resolution fluorescent image taken before photobleaching the region of interest (ROI) where nuclei (cyan), LRP1 (white), and cell borders (dotted lines) are highlighted, b) STORM image and Mean Shift Clustering analysis of the same ROI, with bandwidth at 200 nm c) Violin plot comparison of the clustered number of localization (NOL) and number of molecules (NOM) for LRP1 and single secondary antibody (SA) of Fig. 2.b, d) NOL and NOM clusters distributions of LRP1 and SA at different clustering bandwidths of five different images.

### The monomeric LRP1 Structural Model generation

To build an atomistic structural model of LRP1, we proceeded with the following procedure. As a first step, we evaluate the performance of state-of-the-art structure prediction algorithms like AlphaFold2 [27] and RoseTTAFold [28] predicting the LRP1’s local tertiary structure motifs, shown as examples in Fig. 1.b. The estimated errors made by the two predictive algorithms are reported in Fig. S1.a,b, which assesses the high confidence of the models. Both algorithms successfully produced the three motifs with high agreement among themselves (Fig. 1.c) and with those observed in 8EM4 [18], which is now the largest protein observed in the LDLR family (see Fig. S2). We further evaluated the capacity of AlphaFold3 in producing the full-length LRP1, finding a coiled conformation with many atomic clashes and motif superimpositions (see Fig. S3). The full prediction can not be used for further studies because of its low quality. Hence, we proceeded with predicting portions of the LRP1 sequence with RoseTTAFold as shown in Fig S3. The protein model was obtained by joining the predictions of different domains and further manipulated for better clarity in the representation, following the scheme in Fig. S4. A 3D arrangement of the LRP1 motifs is shown in Fig. 3.a. It is worth to stress that the prediction of each motif is in line with the results of Fig. S1, while the 3D orientation of the motifs was manipulated for obtaining different conformations. After the placement of calcium ions to the calcium binding units’ coordination sites as displayed in Fig. 1.b, the protein was prepared for molecular dynamics simulation as described in the Methods section. The resulting structure is highly flexible, potentially extending to a fully stretched 100nm from the cell surface without altering the tertiary structure of the single domains (as shown in Fig. S4). The latter conformation was obtained by manipulating the dihedral angles to extend the protein but keeping the local motifs intact.

**Figure 3.**
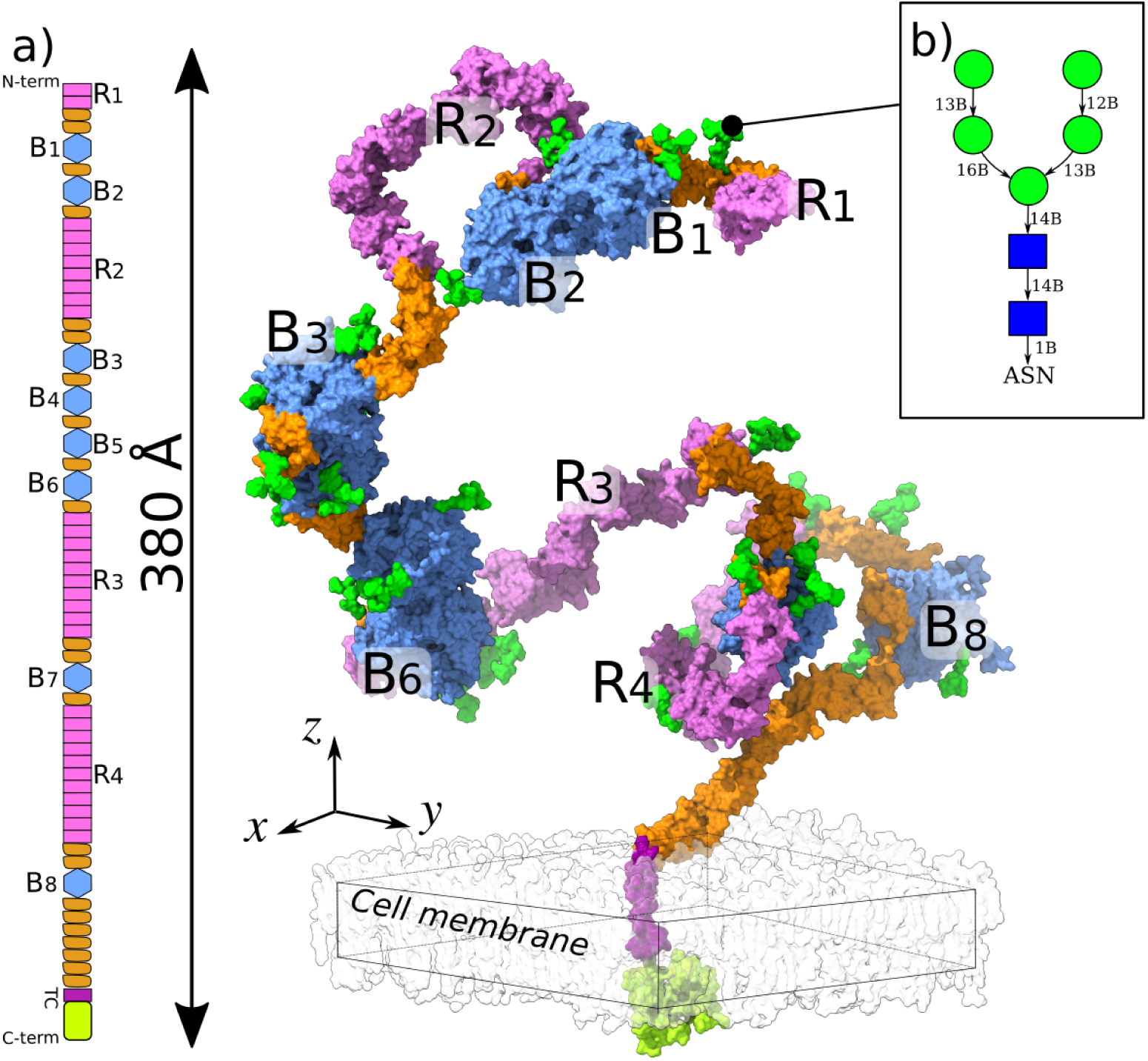
a) The proposed structure of the LRP1 monomer inserted in the cell membrane is represented by a vdW surface. Each local domain follows the colors in (a). Glycans are represented in light green surface. b) 2D representation of the GlcNAc(2)Man(5) glycans used in this work.

### Furin’s shedding side belongs to a **β**-propeller motif

The enzyme furin cleaves LRP1 in correspondence of B8 (Fig. 4.a), generating two chains, α and β, that are non-covalently bonded. To provide information on the shedding mechanism, we investigated the stability of the non-covalent interactions, specifically hydrophobic interactions between adjacent β-sheets (Fig. 4.b,c), which hold the resulting chains together using molecular dynamics (MD) simulations. We isolated the region of ASP3779-GLN4235 from the full monomeric structure of Fig. 3.b and prepared the system as described in the Methods section. We performed a series of short simulations using the umbrella sampling method [29] to estimate the energy required to separate the α and β chains, going from the conformation shown in Fig. 4.b to the final conformation shown in Fig. 3.d. The potential of mean force (PMF) shown in Fig. 4.e, reveals that the energy required to break the interactions between the two chains is approximately 180±2 kT.

**Figure 4.**
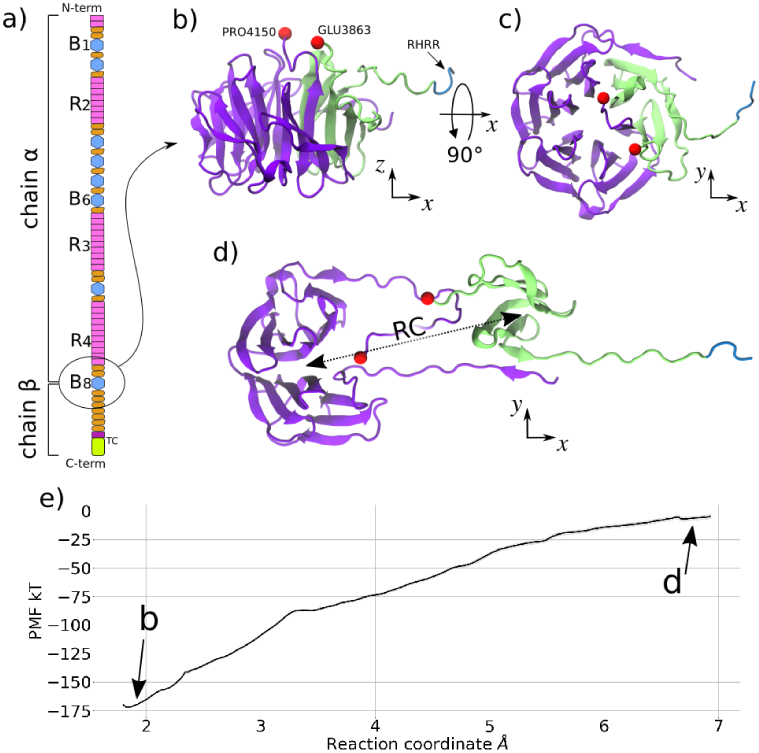
a) The scheme shows the location, along the LRP1 motif sequence, of β-propeller 8 (B8). In correspondence of this motif, the protein is divided into α and β chains. b,c) B8 structure prediction from AlphaFold2, depicted as a cartoon in lime (α chain) and violet (β-chain) with side (b) and top (c) perspective. The RHRR sequence is depicted in blue; the extreme amino acids of the unit are depicted in red spheres. d) B8 at the extreme distance of α and β centers of mass. e) The potential of mean force for the α and β centers of mass distance.

### The dimeric LRP1 Structural Model

To predict LRP1 in a dimeric conformation, we used the crystallized LRP2 structure at pH 7.5 from mice, PDB index 8EM4; the protocol is shown in Fig. 5. As a first step, we modelled the tertiary structure of the mice LRP2 missing amino acids with RoseTTAFold [28]. We confirmed the results with AlphaFold2 [27] prediction. Figure S1 shows the predictive performance of structural prediction algorithms in producing high-quality motifs. The sub-sequences inserted in the RoseTTAFold tool are the sequences of mice LRP2 with residue numbers 1272 to 1349, 2279 to 3033, and 3881 to 3992, hereafter called domains D2, D3 and D4, respectively (in green in Fig. 5.a). The N-terminal of each predicted D-motif was joined to the correspondent site in 8EM4; i.e. the residue 1272 of domain D2 was covalently bonded to the residue 1271 from 8EM4, etc. We ignored the missing residues from LRP2’s R1 since they are not needed for modelling LRP1. We conducted a pairwise alignment of LRP1 and LRP2 sequences using the Needle algorithm [30] (Fig. 5.b). In the LRP2 sequence, we considered the presence of the Dn amino acids. We then ran two homology modelling routines using Modeller [31]. In the first one, Modeller produced the LRP1 ectodomain in the dimeric conformation. Apart from the missing CB units in the flexible R clusters, patched with the D domains, the LRP2 structure also lacked transmembrane and cytoplasmic domains due to limitations of the method. We then predicted these domains with RoseTTAFold and verified with AlphaFold2. Fig. 5.c shows the predicted C-terminal domain colored according to error estimation from RoseTTAFold prediction. The error indicates a good confidence in the alignment of the five EGF-like domains and a lower confidence in the orientation of the transmembrane helix. The transmembrane domain was connected to the C-terminal of the previously obtained dimeric model and oriented to allow insertion into the lipid membrane. Furthermore, a Modeller run added the missing sulphur bridges, constrained the Cys-Cys distances (previously optimised by the RoseTTAFold model), and arranged a homodimer symmetry. Finally, calcium ions were inserted into the calcium-binding sites. The complete structure generated from a homology modelling protocol was visualized in Figs. 5.d and 5.e. The protein complex occupies a 220×280×355 A^°^³ parallelepiped. The protein appears bent in correspondence with the four β-propellers, B3 to B6, which form an arc of 180 degrees. R1, B1, B2, and R2 make the ascending leg; R3, R4, B7, and B8 make the descending leg, which ends up with the transmembrane and cytoplasmic domains.

**Figure 5.**
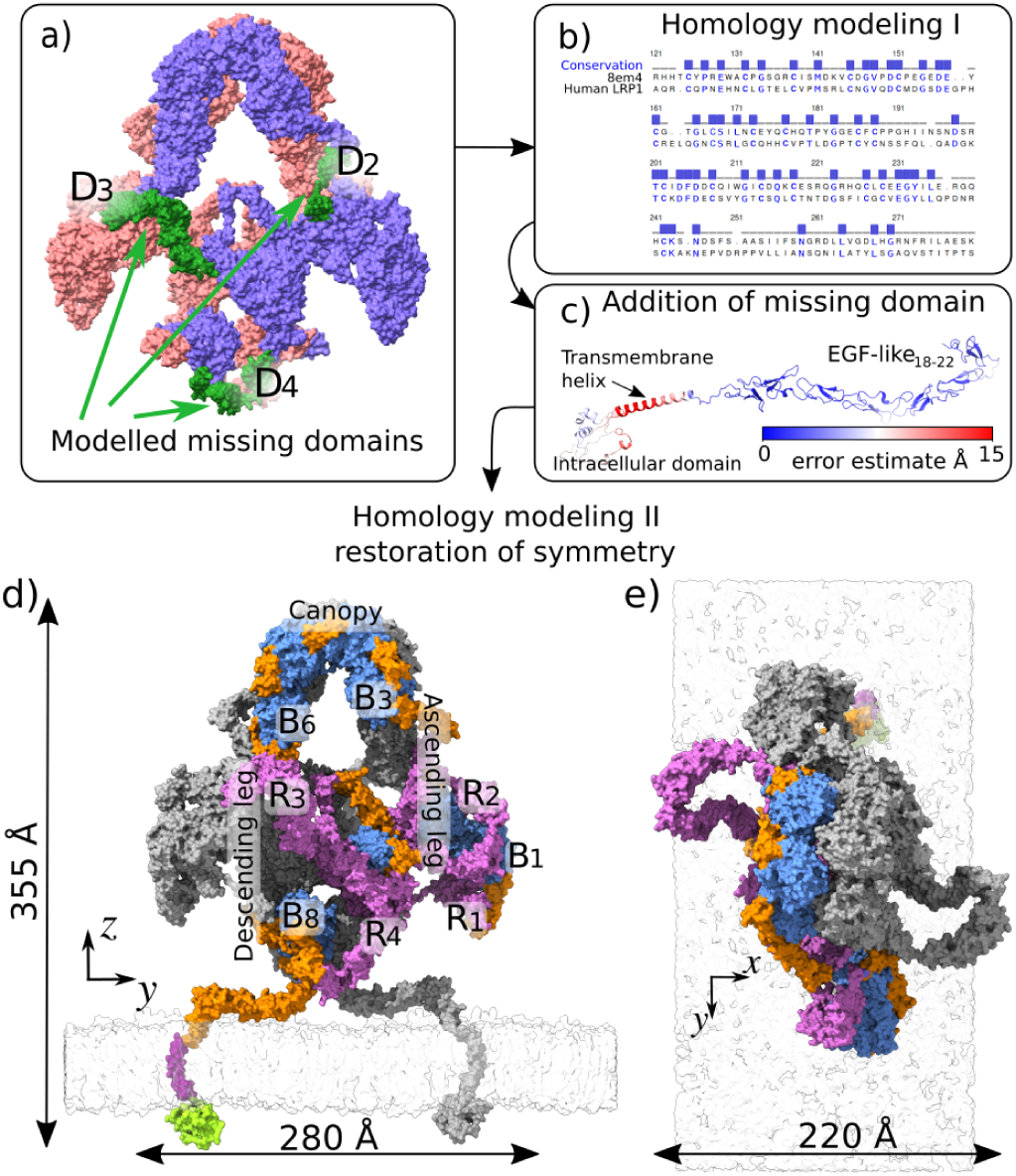
a-c) Protocol steps for the dimeric structure prediction of LRP1. The steps are explained in the text. d) Structure of the LRP1 dimer obtained by homology modelling from PDB 8EM4. The forefront monomer follows the representation in Fig. 1), while the background monomer is in grey. Bn and Rn indicate the β-propeller and the CB domains, respectively. e) Dimer view from z-axis perspective.

The buried interface area between two protomers is 5951 ^°^A², accounting for 1.23% of the protein complex’s surface. The interface area is displayed in Fig. 6.a. The main interaction among protomers occurs in the canopy region. This part of the dimer is to be investigated later in this work. Other intra-dimer interactions are found between the two B8, as shown in Fig. 6.b. As seen in the previous section, the stability of B8 is necessary to keep the LRP1’s α and β chains together. Further intra-dimer contacts could further stabilize the non-covalent interactions between the two monomer’s chains.

**Figure 6.**
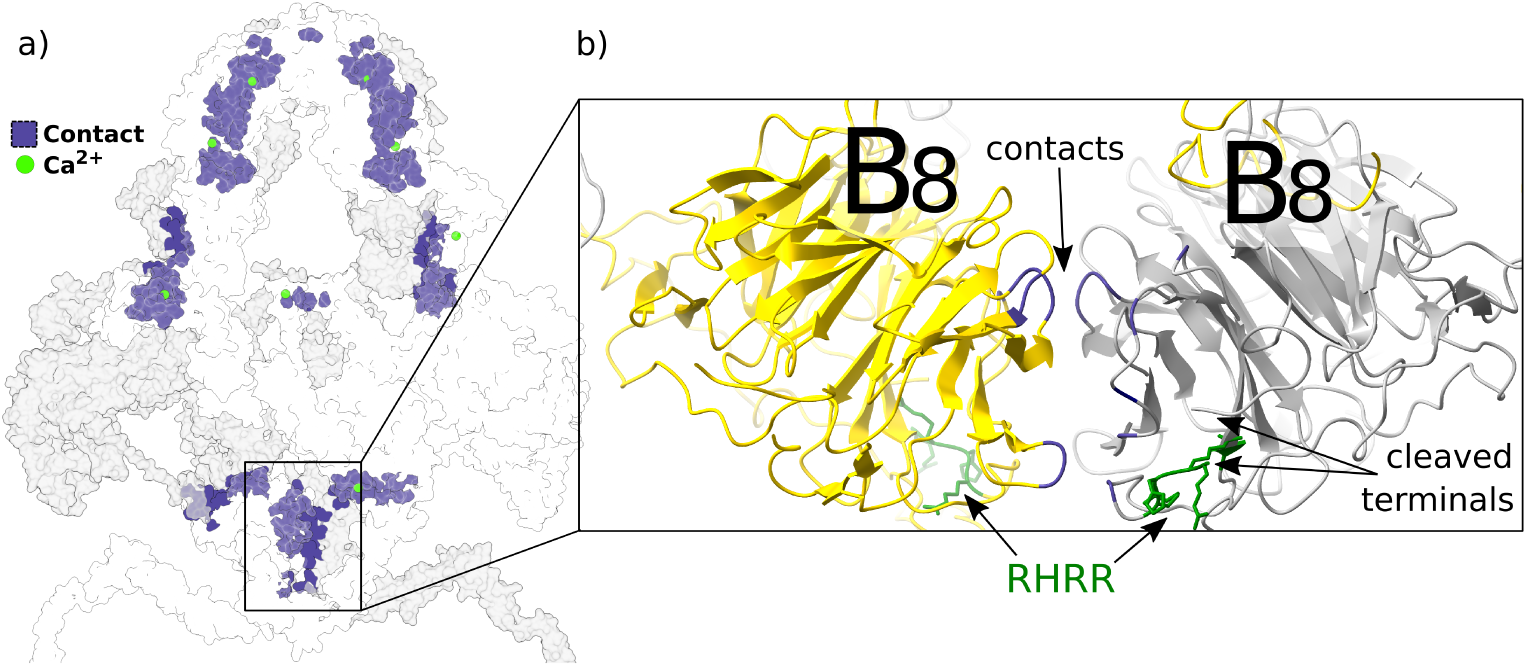
a) Dimeric LRP1 shown as vdW surface; the front monomer is transparent white and the back is light grey. The transparent surfaces allow the vision of hidden contact surfaces (in violet) among the two monomers. b) Zoom in on the lower intra-dimer interface between the two B8, in yellow and grey cartoon, respectively. In green, the two subsequences RHRR after furin cleavage. The interactive amino acids at the interface are colored in violet.

As in the monomer case, the system was prepared for simulations using Charmm-GUI [32], generating the sulfur bridges between Cys pairs along the CB and EGF-like units, and finally, energy minimized (see Methods).

### Time evolution of the flexible domains

The repetition of CB motifs forms three long chains, R2, R3 and R4, as shown in Fig. 1. The flexible behavior of the R domains was observed in the crystal structure of LRP2, where some units of the corresponding clusters have not been resolved with sufficient resolution (Fig. S5), as the single particle analysis identified the domains in different conformations. An interpretation of this fact is that while CB motifs are well structured locally, their relative orientations can vary in time.

We evaluated the temporal conformational changes of the three LRP1 R clusters, isolated by the rest of the protein, with MD simulations. The systems accounted for were: i) R2, amino acids 803 to 1269, CB3 to CB10 and EGF-like4 to EGF-like6; ii) R3, amino acids 2472 to 3025, CB11 to CB20 and EGF-like10 to EGF-like12; iii) R4, amino acids 3289 to 3868, CB21 to CB31 and EGF-like13 to EGF-like15. We decided to start from a stretched conformation to observe the coiling of the domains towards the equilibrium structure. In the process, we did not use any constraints on the domain terminals and observed the system relaxation. The R domains were prepared as discussed in the Methods section. In all of the systems tested, we observed the tendency to coil by measuring the end-to-end distance (d_EE_). We represent the distributions of d_EE_ calculated in the last 200ns of MD simulations in Fig. 7.b. Here, it is visible the reduction of d_EE_ with respect of the initial conformation (dotted black vertical line). Furthermore, we compare the distribution of d_EE_ with the same quantity calculated in the LRP1 dimeric conformation, represented by the red vertical line in Fig. 7.b and shown graphically in Fig. 7.c. We further quantified the flexibility of the three R domains by calculating the root mean square fluctuation (RMSF) of the alpha-carbon of each amino acid with respect to the initial positions. In Figure 6.d, we show the initial and final conformations of each domain, with the latter colored with respect to the RMSF.

**Figure 7.**
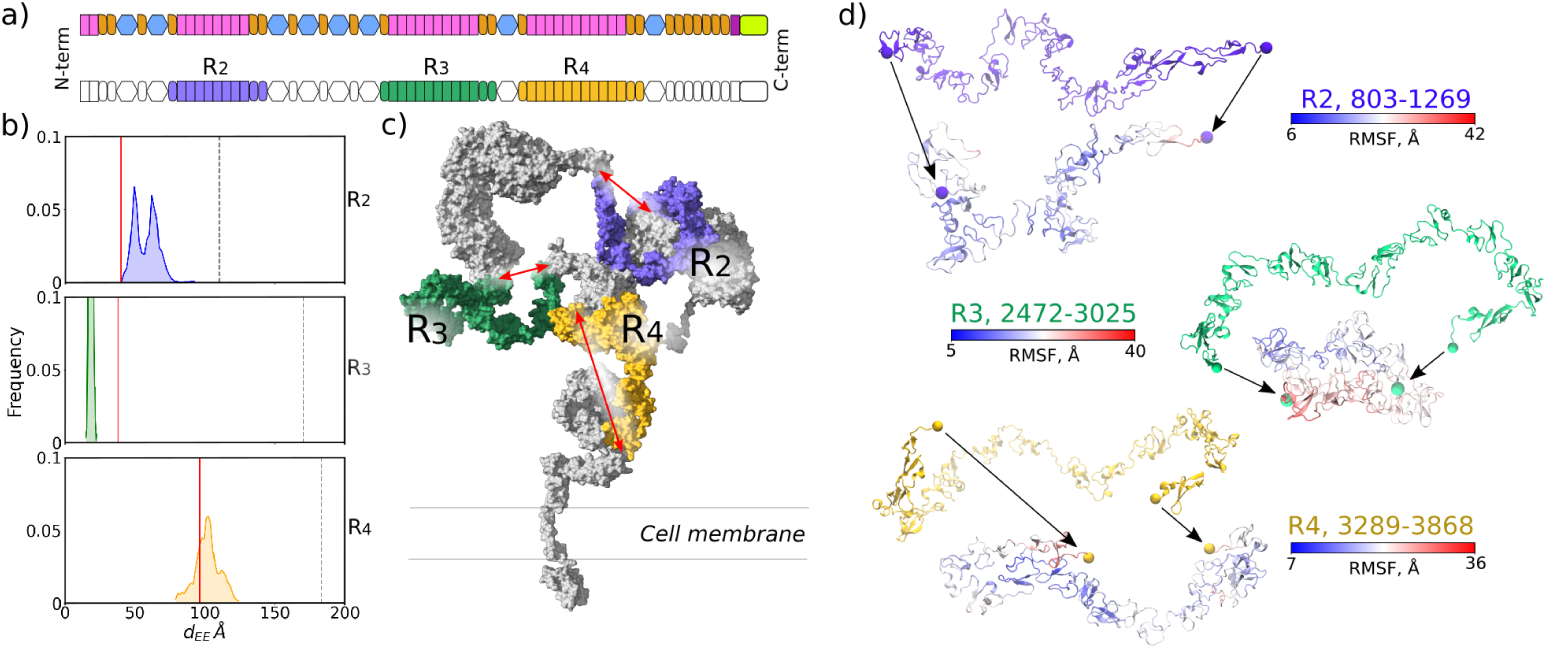
a) Schemes of LRP1 motif sequence. Top: same scheme as Fig. 1. Bottom: Same scheme, enhancing with colors the motifs of macrodomains R2, R3, and R4. b) Normalized frequency distribution of the Rn domain end-to-end distance dEE during the last 200 ns of MD simulations. The vertical dotted line represents the dEE value in the initial conformation; the red solid line is the dEE value for the R domain in the dimeric conformation. c) Visual representation of LRP1 monomer in vdW surface. The three domains, R2, R3, and R4, define a static end-to-end distance, highlighted by the red arrows. d) Initial and final conformations of the R domains. The initial conformation is depicted following the color scheme; the final is colored according to the calculated RMSF of each amino acid with respect to the initial conformation. The black arrows follow the translation of the domain terminals, depicted with vdW spheres.

During the evolution of the flexible domains, we did not observe any divalent calcium ions leaving their respective coordination sites. We investigated the strength of the interaction between the ion and its coordination site by employing umbrella sampling and the Weighted Histogram Analysis Method (WHAM) [29] (see Methods). For the procedure, we choose to use the CB7 from R2, which displays the two structural features of CBs: i) the conservation of all six Cys; ii) the conservation of the acidic amino acids in the calcium cage: Asp1035, Asp1037, Asp1039, Asp1045 and Glu1046 (Fig. 8.a). The results of umbrella sampling (Fig. 8.b) show a potential of mean force (PMF) with a deep equilibrium well. As the reaction coordinate increases, the system finds other equilibrium conformations that are less stable with respect to the original one. In this transient state, the ion is geometrically outside the cage. Finally, the ion is free from the cage interactions. The free ionic conformation is estimated to be reached after giving the system 16±2kT of energy, justifying the rarity of this event.

**Figure 8.**
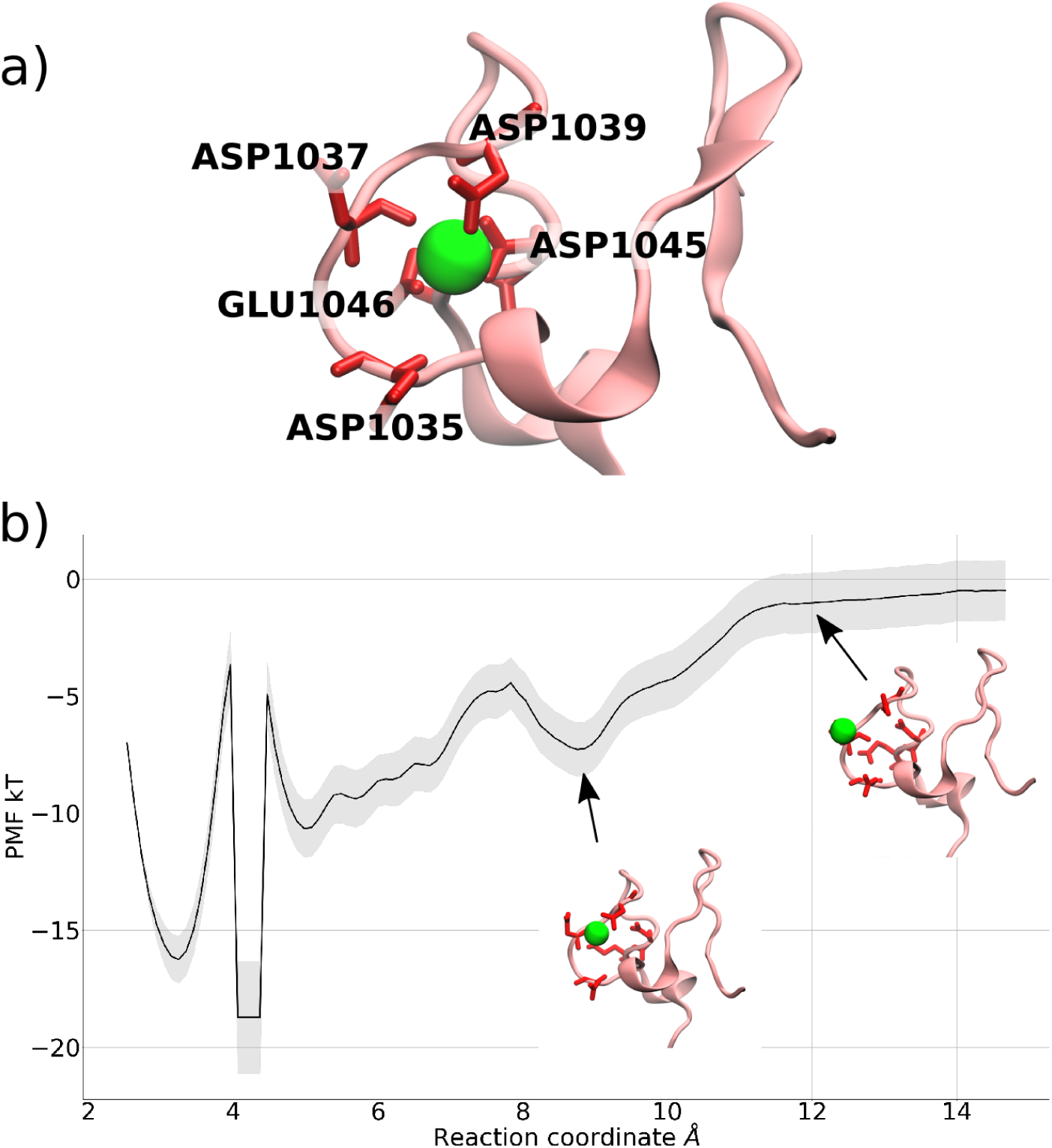
a) CB 7 from R2 displayed as pink cartoon. Key amino acids are shown as red sticks. The calcium ion is a green sphere. b) Energy profile from the umbrella sampling protocol. The shaded area shows the uncertainty of the measure. The inset cartoons show local configurations of the ion.

### The protein glycosylation affects the LRP1 dimer conformation

An interesting feature of LRP2 and LRP1 dimers is the large interaction surface between the two monomers in correspondence with the canopy area (Fig. 9.a). Since this area is densely glycosylated, we investigated the action of N-glycans on the stability of the dimeric interaction. The analysis focused on amino acids 1140 to 2522 of both monomers, as shown in Fig. 9.a. This fragment of the full LRP1 contains β-propellers B3, B4, B5 and B6, as well as the EGF-like5-10 motifs. All the glycans are N-type and bonded to the corresponding glycosylation sites reported in the GlyGen database [33]. This repository assesses the presence of twenty-one glycosylation sites for each monomer in the considered area. We decided to include only 19 of them, ignoring the ones corresponding to ASN1155, which raises steric hindrance problems with another glycan on ASN1154, and ASN2521, the system’s second-to-last amino acid. The final count of glycans resulted in 38 glycosylation sites. The glycans are the same simple high-mannose used for the flexible domains, as shown in Fig. 3.c. The sugar chains were added with the Glycosylator software [34], and they needed to be further manipulated to avoid clashes in the small space between the protein canopies. The system, glycosylated and with four calcium ions at the interface (as described by Beenken and colleagues [18]) was prepared for MD simulations with the Charmm-GUI webserver [32].

**Figure 9.**
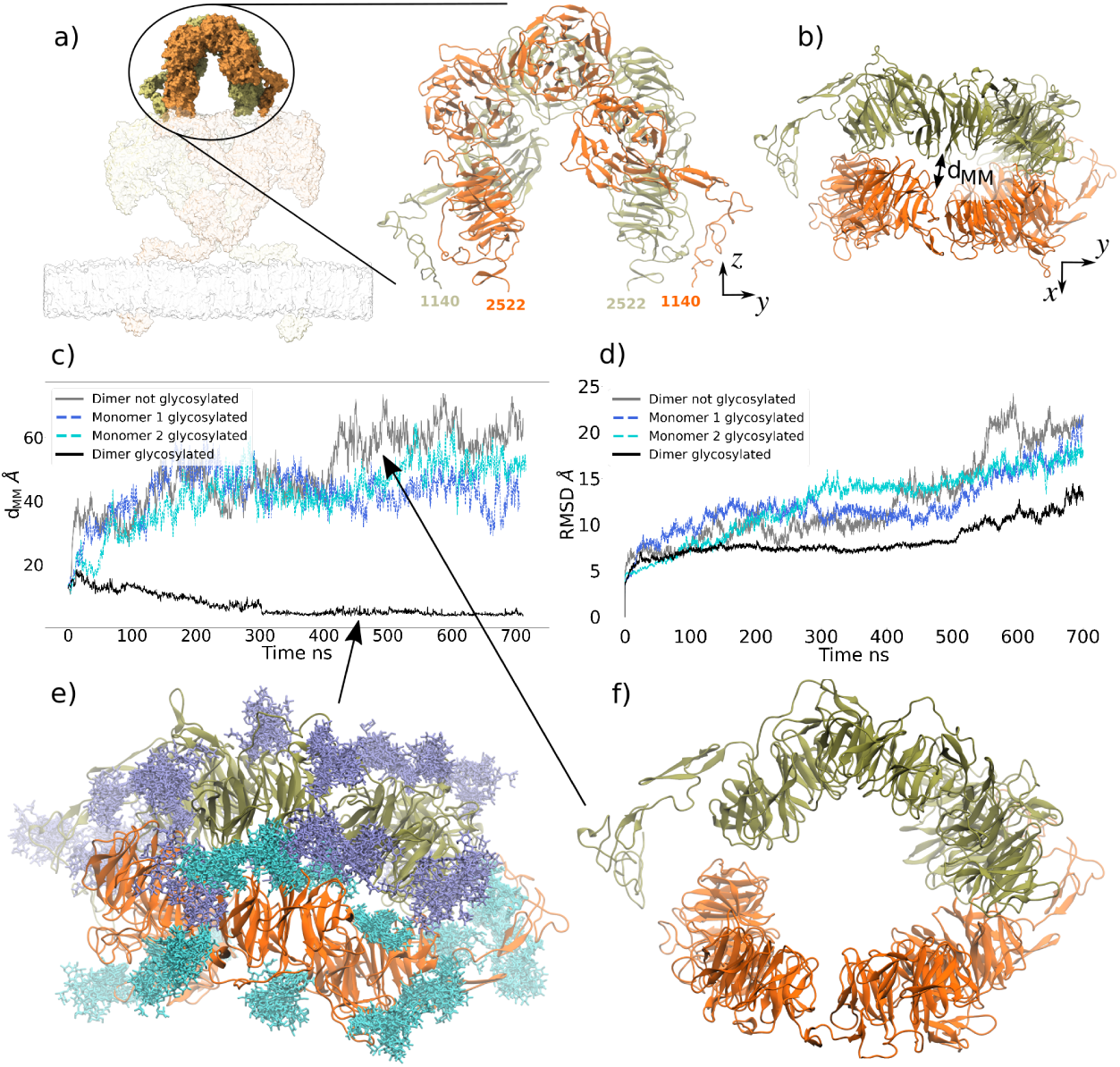
a) Detail of the dimeric canopy structure. The two monomers are colored orange and khaki. b) The xy view of the dimeric canopy highlights the protein vicinity, quantified by the distance d_MM_. c) d_MM_ in function of time for the four systems: glycosylated dimer, asymmetrically glycosylated monomers, and non-glycosylated. d) Root Mean Square Distance of heavy atoms for the four systems as a function of time. e) Glycosylated canopy structure after 500ns; ice blue and cyan represent the N-glycans bonded to the two proteins. Ten conformations are shown for each glycan, highlighting the space occupied during the system evolution. The proteins keep their initial distance. f) Non-glycosylated canopy structure after 500ns. The proteins are mainly detached, creating a pore-like structure.

We conducted four sets of classical MD simulations of the dimeric canopy region: i) fully glycosylated monomers, ii)-iii) asymmetric glycosylation of only one dimer member; iv) no glycosylation. Systems were let to evolve for 500ns using constraints on the free terminals, reproducing the effect of the inertial mass of the rest of the protein. After the first 500ns, we relaxed the positional constraints on the terminals and let the systems equilibrate for another 200ns. The changes in protein conformations are determined by the monomer-monomer distance between two amino acids, namely d_MM_, illustrated in Fig. 9.b and reported as a function of time for all the systems in Fig. 9.c. This tendency is further confirmed by the dimeric RMSD shown in Fig. 9.d.

We observed a discrepancy in the systems’ behavior. The fully glycosylated canopy keeps the initial conformation, called hereafter “closed”. In contrast, the partially glycosylated and non-glycosylated system undergoes a conformational change at the beginning of simulations, assuming an “open” conformation that is kept throughout the simulation. The final conformations for the closed, fully glycosylated, and open, non-glycosylated canopies are shown in Figs. 9.e and 9.f, respectively.

To quantify the conformational difference between the closed, fully glycosylated system and the open, non-glycosylated system, we investigated the hydrogen bonds (HB) between the systems’ entities. We identified the possible HB interactions as: i) protein-protein; ii) protein-glycan; iii) glycan-glycan (Fig. 10.a). The fully glycosylated presented all three, while the non-glycosylated presented only protein-protein interactions. In Fig. 10.b we show the distribution of HB for the two systems, calculated in the range 400-500ns (with constrained terminals) and 600-700ns (after relaxation). After relaxation, we observed an increase in interactions between the monomers for both the open and closed conformations. The non-glycosylated configuration exhibited an average increment of three HB per frame, located among B3 and B6, which stabilized the open conformation.

**Figure 10.**
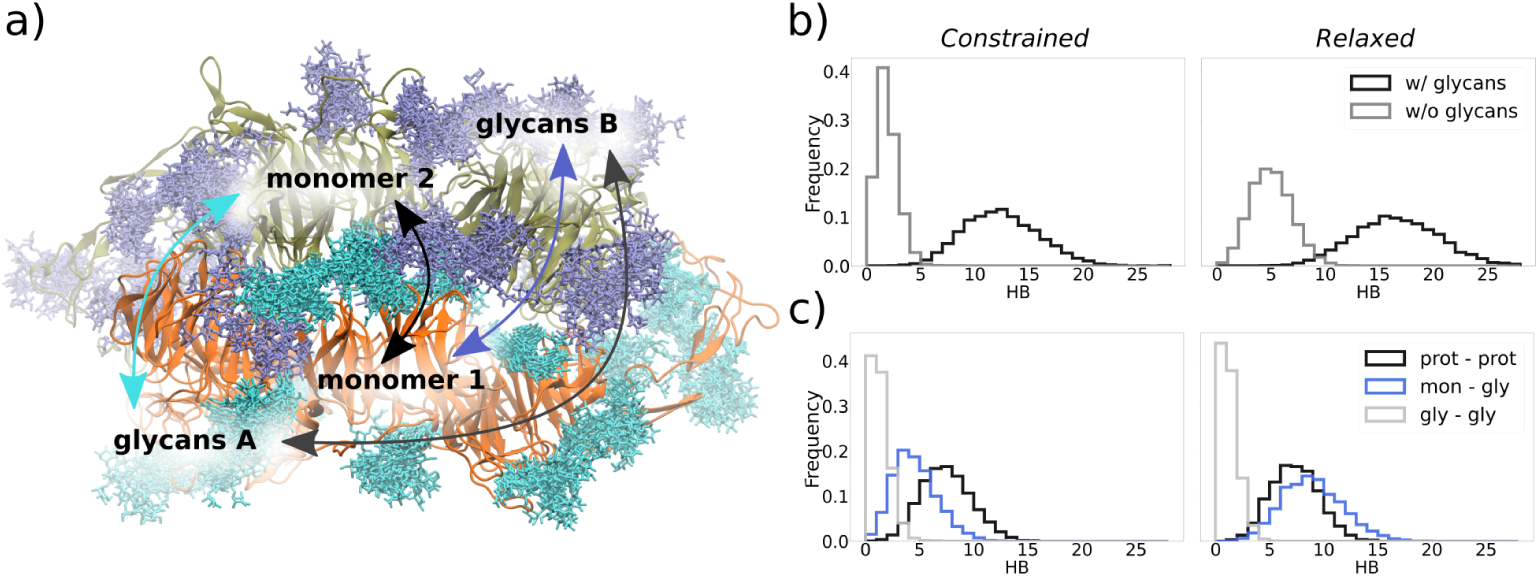
a) Detail of the dimeric canopy structure in closed conformation at the end of the simulated time (700ns). The monomers 1 and 2, are colored orange and khaki, respectively; the two glycan groups, A and B, are represented as licorice and anchored to monomer A and B, respectively. b) The distribution of hydrogen bonds was calculated in the constrained and relaxed set up for the closed and open conformations. In the closed conformation, fully glycosylated, the protein-protein, protein-glycan and glycan-glycan components were summed. In the open conformation, only the protein-protein HB was calculated. c) The HB distribution for the three components in the fully glycosylated system: protein-protein, protein-glycan and glycan-glycan interactions. The second is the sum of monomer 1 – glycan B and monomer 2 – glycan A contributions.

With respect to the fully glycosylated system, an average increase of four HB was visible. In Fig. 10.c, we show the HB components for the fully glycosylated system. While the protein-protein HB displayed a slight depletion, reaching equilibrium in a distribution comparable with the open conformation, the number of protein-glycan interactions notably increased. The number of glycan-glycan interactions was small and did not significantly influence the system’s time evolution.

Notably, the non-glycosylated system tended to have less stable hydrogen bonds during the 700ns simulated time (Tab. 1).

**Table 1.** First four intra-dimer hydrogen bonds, ordered by stability, for the glycosylated and non-glycosylated canopy, before (400-500ns) and after (600-700ns) the relaxation of terminal constraints. For each amino acid couple, we listed the ConSurf score.

| Glycosylated |  |  | Glycosylated - relaxed |  |  |
| --- | --- | --- | --- | --- | --- |
| HB | Occ % | ConSurf score | HB | Occ % | ConSurf score |
| ARG1497<br>GLU2418 | 93.2 | 8-5 | ARG1966<br>ASP1407 | 87.6 | 7-9 |
| ARG1517<br>GLU2418 | 87.9 | 7-5 | TYR2030<br>PRO1411 | 61.5 | 3-9 |
| TYR2030<br>PRO1411 | 61.8 | 3-9 | TYR2030<br>SER1409 | 54.9 | 3-6 |
| TYR2030<br>SER1409 | 57.1 | 3-6 | LYS1788<br>GLU1899 | 53.3 | 4-6 |
| Not glycosylated |  |  | Not glycosylated - relaxed |  |  |
| HB | Occ % | ConSurf score | HB | Occ % | ConSurf score |
| ARG1517<br>GLU2418 | 39.5 | 7-5 | ARG1517<br>GLU2418 | 59.1 | 7-5 |
| ARG1517<br>ASP2399 | 21.2 | 7-9 | ARG1517<br>ASP2399 | 41.6 | 7-9 |
| LYS2454<br>GLU1476 | 19.6 | 6-8 | LYS2416<br>PHE1477 | 30.2 | 5-5 |
| ARG1517<br>PRO2419 | 15.2 | 7-8 | ARG1454<br>ASP2008 | 29.9 | 7-8 |

We further assessed sequence conservation of LRP1 using ConSurf [35] (Fig. S2) (see Methods). It was identified that amino acids located in the core of β-propellers were highly conserved (Fig S2.a and d), whilst residues located on the surface were generally less conserved, apart from residues located at the dimer interface (Fig S2.b and c). Conservation of residues at the dimer interface highlighted their potential role in maintaining protein-protein interactions. However, by comparing evolutionary data with molecular dynamics results, we identified that amino acids involved in intra-dimer HBs in the non-glycosylated system had higher conservation scores but formed less stable HBs than the ones in the glycosylated system. The comparison suggested that conserved amino acids at the dimer interface are less relevant than glycans to the maintenance of the dimer structure.

## Discussion

We present the novel hypothesis of a multi-quaternary structure for LRP1. The STORM analysis highlights the possibility for LRP1 to cluster on the surface of brain endothelial cells. With the available data, it is not possible to affirm what interaction exists between two or more LRP1, nor if it is direct or mediated by other entities; however, the observation of groups of LRP1 indicates the presence of an interaction. After this evidence, we produced two novel models for the LRP1, showing both the monomer and homodimer structure.

In the absence of significant experimental details on the tertiary structure, the monomeric structure of LRP1 is presented as a multi-domain flexible protein inserted into the cell membrane. Flexibility highly depends on the three calcium-binding domains R2, R3, and R4.

LRP1 is shed into two chains, α and β, by furin in the trans-Golgi compartment [10]. The shedding site appears in the β-propeller 8 (B8) motif. We investigated the stability of the B8 motif. We quantified the energy necessary to break the non-covalent bonds between the two chains, finding it to be around 180±2 kBT and hence impossible to break without a specific event. This finding agrees with the literature of soluble LRP1 (sLRP1), a species of LRP1 shed from the membrane of cells under stress [36, 37]. Even in blood circulation, under application of strong torque forces, the non-covalent bond of α and β chains is conserved in sLRP1.

The homodimer model was obtained using a combination of neural network-based models for protein structure prediction and traditional homology modelling from the structure of a close homologous receptor, LRP2. The obtained structure keeps the same features as the template: a quaternary structure, intra-dimer contacts of rigid domains, and more flexible domains free of evolving in water. Moreover, we advance a hypothesis for the positions of transmembrane and intracellular domains that are not present in the LRP2 template. The dimerization of LRP1 was never observed in physiological experiments. The novelty of this result is of great biological importance: if LRP1, like LRP2, can form homodimers, it raises the possibility of forming heterodimers with the other members of the LDL receptor family, giving birth to unique pathways.

The two structures raise the possibility of analyzing LRP1 through different approaches, including MD simulations. Our first analysis studied the time evolution of flexible domains R2, R3, and R4. The three domains were isolated from the rest of the protein and artificially elongated while keeping all the secondary structures intact. These domains show a tendency to coil. The final conformations of R2, R3, and R4 show, at equilibrium, an end-to-end distance comparable to the same domains in the homodimer structure, as reported in Fig. 7. This fact indicates the thermodynamic equilibrium of the R domains in the proposed dimeric conformer, validating the latter. We did not observe a change in the coiling behavior influenced by the presence of glycans.

We observed calcium stability in the R domains’ coordination sites. Not a single calcium ion left its site during the MD simulations. We investigated the strength of this bond, finding it is due to two factors: the opposite charges of Ca^2+^ and its cage, and the geometric distribution of the cage amino acids surrounding the ion. We employed umbrella sampling to force the ion outside the cage (Fig. 8), finding that the PMF difference between the bound and unbound state is 16±2 kT. We hypothesize that the energy barrier is even higher; however, we were unable to quantify it.

We conducted a study on the canopy region of the LRP1 homodimer. This structure comprises four β-propellers motifs for each protein, disposed in a “U” shape. The two proteins interact laterally through β-propeller interactions. As this region is believed to be highly glycosylated, we tested the stability of the native interactions in the absence and presence of glycans. The presence of glycans notably increases the amount of intra-dimer interactions, making the quaternary structure more stable in the closed dimer conformation. Without glycans, the hydrogen bonds between the two proteins are less stable. Thus, the open structure is not as solid as the closed one.

Evolutionary sequence analysis of the canopy β-propellers (see Fig. S2) indicated high conservation of residues in comparison with the rest of the protein, with core residues being more evolutionarily conserved than those at the surface. Conservation of core residues matched previously described YWTD motifs [15]. Further differences of canopy β-propellers were observed with β-propellers 3 and 6 having the most conserved residues, highlighting a further interesting evolutionary pattern of the motifs. A certain glycosylation pattern may promote a closed conformation and compensate for the divergent mutations at the surface of β-propellers 5 and 6, which instead promote an open conformation of LRP1.

A dynamic analysis of the intra-dimer hydrogen bonds indicated weak interactions for more conserved amino acids in the absence of glycans, and strong interactions for less conserved amino acids in the presence of glycans, suggesting a non-functional role of conserved amino acids of the canopy domain in LRP1 dimerization.

Together with our other observations, we conclude that glycans could have a fundamental role in allowing dimerization and influencing the quaternary structure conformation of LRP1 and, potentially, of different proteins.

## Materials and Methods

### Immunofluorescence staining of bEnd.3

Mouse brain endothelial cells (bEnd.3) were seeded onto fibronectin-coated µ-Slide 18-well Glass Bottom chambers (IBIDI) and cultured to confluence over 3 days. Cells were rinsed with PBS and fixed in 3.75% (w/v) paraformaldehyde for 10 minutes at room temperature. To preserve surface-exposed LRP1, no permeabilization was performed. Fixed cells were blocked with 5% bovine serum albumin (BSA) in PBS for 1 hour at room temperature and incubated overnight at 4°C with mouse anti-LRP1 monoclonal antibody (Huabio, M1211-4) diluted 1:50 in 1% BSA in PBS. The following day, after three washes with PBS, cells were incubated for 1 hour at room temperature with donkey anti-mouse IgG2a Alexa Fluor 647-conjugated secondary antibody (Abcam, ab150107) diluted 1:200 in 1% BSA in PBS. Nuclei were counterstained with Hoechst 33342 (1:1000 in PBS), followed by three final PBS washes. Samples were imaged immediately.

### STORM imaging set-up and secondary antibody calibration

To achieve the fundamental principle behind stochastic optical reconstruction microscopy (STORM), cells were incubated during the imaging with a GLOX buffer containing (i) 2 µL of GLOX, (ii) 20 µL of MEA, and (iii) 20 µL of glucose (50% w/v) diluted in 160 µL of PBS pH 7.4. This GLOX buffer is the most suitable for 647-conjugated secondary antibodies, as it is based on an oxygen scavenging system that oxidizes Alexa Fluor 647 and reduces its blinking properties. We also calibrated the secondary antibody to identify the number of localizations produced by a single molecule. Therefore, the goat anti-mouse IgG2a 647-conjugated secondary antibody was diluted 400,000 times in PBS and placed on a glass coverslip (Corning Cover Glass, thickness 1½, 22 × 22 mm) for 10 minutes. Several washes with PBS were performed to remove the excess antibody. The same GLOX buffer was used to facilitate the imaging of the secondary antibody.

### STORM imaging and Mean Shift Clustering

STORM imaged LRP1 receptors and the secondary antibody calibration by illuminating the sample with the 647 nm (160 mW) laser at 50% power for the first 10,000 frames and then at 100% for the successive 10,000 frames. Seven images were taken for LRP1 samples, while two were for secondary antibody calibration. Fluorescence was collected using a Nikon 100× oil immersion objective with a numerical aperture of 1.49. The light passed through a quad-band pass dichroic filter (97335 Nikon). Images were captured over a 256 × 256 pixel region, with a pixel size of 0.16 µm, using a Hamamatsu ORCA Flash 4.0 camera at an integration time of 10 ms. STORM movies were analyzed with Nikon’s NIS Elements software, employing a 2D Gaussian fitting with thresholds of 200 and 250. The threshold represents the distinction between the photon counts in the peak versus the background pixels. The trace length parameter was set to 5 to prevent the consecutive blinking of different molecules. This means that if a molecule is seen in five consecutive frames, it counts as one; if it appears in more than five frames, it will be removed. Signals with more than five localizations have been considered valid for molecular quantification. Then, we carried out the Mean Shift Clustering (MSC) analysis to identify clusters and evaluate the number of localizations. The MSC was performed by selecting arbitrary radii of 50, 100, and 200 nanometers. Outliers were removed from the clusters identified by Mean Shift Clustering by the ROUT method (Q=1). Afterwards, descriptive statistics of cluster distributions were made to determine the mean, median, and percentiles.

### LRP1 Monomer Modeling

The monomeric structure of LRP1 was formulated based on the knowledge of the sequence of protein structural motifs [16]. After assessing the validity of RosettaFold’s structural predictions (Fig. S1), we predicted long portions of LRP1 (see Fig. S4).The so-obtained protein portions were combined in Chimera [38]. The calcium ions were added to the coordination site of the CB motifs, as shown in Fig. 0.b and observed in the members of the LDL receptor family [12, 18]. We then generated a structural ensemble for the monomeric unit using classical molecular dynamics simulations. The system was prepared using the Charmm-GUI web server [32], generating the sulfur bridges between Cys pairs along the CB and EGF-like units, and adding the N-glycans. The glycosylation of proteins, that is, the covalent attachment of carbohydrates to proteins, is a posttranslational process highly dependent on many factors and dictated by the necessities of tissues and single cells [39]. Furthermore, the human glycans are the most complex; it is likely that LRP1 presents many different types of glycosylations at different glycosylation points [40, 41]. We decided to decorate the LRP1 monomer’s 52 N-glycosylation points with the same high-mannose sugar chain GlcNAc(2)Man(5) (Fig. 3.b). This glycan represents the sugar’s steric volume. Finally, we conducted an energy minimization without the membrane bilayer (see next Methods section).

### Molecular Dynamics Simulations

#### Full single protein and dimer minimization

The complete LRP1 structures were minimized in OpenMM 8.0 [42] using the Langevin Integrator [43] to a tolerance of 1kJ/mol. The force fields used are AMBER ff14SB for protein, GLYCAM06j-1 for glycans, and standard TIP3P [44] calcium ions. Since the dimensions of the systems are prohibitively large, we opted for energy minimization in implicit solvent, using the Generalized Born-Neck2 solvation method [45]. As the energy optimization was carried on without periodic conditions, the non-bonded interaction cut-off was set to 1nm. The repartition of hydrogen mass was not used.

#### Estimation of the energy required to separate the **α** and **β** chains using umbrella sampling

The structure of LRP1 B8, shown in Fig S12, was prepared for the simulations with Charmm-GUI [32]. The system was solvated with TIP3P [44] water and KCl 0.15 M in a cubic box of 14.9nm side. All the MD simulations were carried on with Gromacs 2021.2 [46], using Charmm36m as protein force field [47]. The system was minimized with the steepest descent algorithm with a force tolerance of 1000 kJ/mol/nm and constraints on protein backbone and sidechains of 400 and 40 kJ/mol, respectively; 4kJ/mol dihedral restraints were applied on glycans. We then proceeded with the first NVT 125ps equilibration step. The algorithm LINCS manages the constraints. We used the velocity rescale thermostat with 1ps coupling constant, which keeps the system heated at 303.15K. Van der Waals force has a cut-off at 1.2 nm, and force-switch is activated at 1.0 nm. Coulomb force is applied with the fast smooth particle-mesh Ewals (PME) method; the cut-off is fixed at 1.2 nm. The NVT productions were carried on with the same parameters.

After the initial equilibration, we generated the initial configurations for umbrella sampling. The protein chains were pulled apart with a constant force of intensity 300 kJ mol^-1^ nm^-2^ during 25 ns with timestep 2fs and a rate of change of the reference position of 10nm per ns. Van der Waals long range corrections were applied for energy and pressure. The chosen ensemble was NPT, using Parrinello-Rahman barostat with coupling constant of 0.5 ps, compressibility of 4.5×10^-5^ bar^-1^ and target pressure of 1 bar.We selected 26 configurations on which to apply umbrella sampling, at intervals of 2 A^°^ with respect to the distance of α and β centers of mass (only amino acids in motif B8, ASP3779-GLN4235). Every configuration was equilibrated for 100ps using the previous step’s setup. Then, we run 10ns umbrella sampling productions using the previous step’s setup. The umbrella sampling analysis used the Gromacs package Weighted Histogram Analysis Method (WHAM). The error of the method was estimated using the Bayesian Bootstrap analysis [48], conducting 200 bootstraps.

#### Dynamics of the flexible domains

All the fragment simulations were carried out in Gromacs 2021.4 [46]. The systems were produced with Charmm-GUI [32]. The systems were solvated with TIP3P water [44] in cubic boxes and implemented with the Charmm36m force field [47]. Systems were minimized with the steepest descent algorithm with a force tolerance of 1000 kJ/mol/nm and constraints on protein backbone and sidechains of 400 and 40 kJ/mol, respectively. The systems containing glycans were also subject to 4kJ/mol dihedral restraints. We then proceeded with the first NVT 10ns equilibration step. The algorithm LINCS manages the constraints. We used the velocity rescale thermostat with 1ps coupling constant, which keeps the system heated at 303.15K. Van der Waals force has a cut-off at 1.2 nm, and force-switch is activated at 1.0 nm. Coulomb force is applied with the fast smooth particle-mesh Ewals (PME) method; the cut-off is fixed at 1.2 nm. These settings are used for all the calculations. During the NVT equilibration, conducted with timestep 1fs, we started with the same constraints as the ones used in the minimization step. We then halved the constraints after 2.5ns and 5ns. The last 2.5 ns were conducted without constraints. After the systems had been thermalized, we proceeded with a 50ns NPT equilibration, using the Parrinello-Rahman isotropic barostat with coupling constant 5ps, target 1 bar and compressibility 4.5×10^-5^ bar^-1^. During this step, we increased the timestep to 2fs. We proceeded with NVT production steps of 100ns until equilibrium was reached. The MD trajectories were analyzed with the MDAnalysis python package [49]. The end-to-end distance dEE describing the coiling of the domains was calculated with respect to the alpha-carbon of the amino acids 852-1182, 2522-2940 and 3332-3778 for macrodomains R2, R3 and R4 respectively. These terminals were chosen to consider only the fluctuations of the CB motif sequences, ignoring the motion of the EGF-like units. The same quantity was measured from the dimeric structure for comparison. The root mean square fluctuation (RMSF) per amino acid was calculated with VMD [50]. We extracted from the trajectory one conformation every 100ns (eight frames in total) and rigidly aligned them to the initial conformation minimizing the root mean square deviation for each atom. Finally, we used the Timeline RMSF calculator setting window width to 8 and step size to 1.

#### Extraction of calcium from its coordination site using umbrella sampling

The CB7, isolated from the full protein model of Fig. 3.b and its Ca^2+^, was prepared for the simulations with Charmm-GUI [32]. The system was solvated with TIP3P [44] water and KCl 0.15 M in a cubic box of 15.6nm side. All the MD simulations were carried on with Gromacs 2020.6 [46], using Charmm36m as protein force field [47]. After a steepest descent minimization, the cube was subject to 500ps NPT equilibration using Nose-Hoover thermostat (reference temperature of 303.15 K and time coupling constant of 1ps) and isotropic Parrinello-Rahman barostat (reference pressure of 1 bar, time coupling constant of 5ps and compressibility of 4.5e^-5^ bar^-1^) and timestep 1fs. The algorithm LINCS manages the constraints. Van der Waals force has a cut-off at 1.2 nm, and force-switch is activated at 1.0 nm. Coulomb force is applied with the fast smooth particle-mesh Ewals (PME) method; the cut-off is fixed at 1.2 nm.

After the initial equilibration, we generated the initial configurations for umbrella sampling. The protein and the calcium ion were pulled apart with a harmonic potential of constant 1000 kJ mol^-1^ nm^-2^ during 2 ns with timestep 2fs, coupling constant of 1ps and rate of change of the reference position of 10nm per ns. The protein backbone was constrained with a force of 400 kJ mol^-1^ nm^-2^. Cut-off for non-bonded interactions were set to 1.4 nm. Van der Waals long range corrections were applied for energy and pressure. The chosen ensemble was NVT, with reference temperature 303.15K. We selected 25 configurations on which to apply umbrella sampling, at intervals of 2 A^°^ with respect to the distance of protein and calcium centers of mass. Every configuration was equilibrated for 100ps using the previous step’s setup. Then, we run 10ns umbrella sampling productions using the previous step’s setup. The umbrella sampling analysis was conducted using the Gromacs package Weighted Histogram Analysis Method (WHAM). The error of the method was estimated using the Bayesian Bootstrap analysis [48], conducing 200 bootstraps.

#### Dynamics of the dimeric interactions

All the canopy simulations were carried out in Gromacs 2022.4 [46]. The systems were produced with CharmmGUI [32] after adding glycans to the protein portions with Glycosylator [34]. The systems were solvated with TIP3P water [44] in cubic boxes and implemented with the Charmm36m force field [47] and multi-site Ca^2+^ ions [51]; then were minimized with the steepest descent algorithm with a force tolerance of 1000 kJ/mol/nm. We proceeded with five NVT equilibration steps of 1ns each, raising temperatures 10, 50, 100, 200, 300. The algorithm LINCS manages the constraints. We used the Nose-Hoover thermostat with 1ps coupling constant, which keeps the system heated at 303.15K. Van der Waals force has a cut-off at 1.2 nm, and force-switch is activated at 1.0 nm. Coulomb force is applied with the fast smooth particle-mesh Ewals (PME) method; the cut-off is fixed at 1.2 nm. These settings are used for all the calculations. During the NVT equilibration, conducted with timestep 1fs, we applied constraints on the proteins’ terminals and Ca^2+^ ions. After the systems had been thermalized, we proceeded with a 1ns NPT equilibration, using the Parrinello-Rahman isotropic barostat with coupling constant 5ps, target 1 bar and compressibility 4.5×10^-5^ bar^-1^. We proceeded with NVT production of 500ns and timestep 2fs, consistently applying spatial constraints on the proteins’ terminals. Finally, we removed the constraints and ran another production of 200ns. The MD trajectories were analyzed with the MDAnalysis python package [49].

#### Evolutionary sequence analysis

A query sequence of the full human LRP1 sequence (Uniprot ID: Q07954) was used as an input for the ConSurf server (https://consurf.tau.ac.il/) [35] to estimate sequence conservation. Sequence homologue search was performed using a single HMMER iteration against Uniref-90 database with E-value cutoff of 0.0001. A set of 150 homologue sequences with minimal and maximal percent identity of 35 and 95, respectively, to the reference sequence were selected for analysis. Selected sequences were aligned using MAFFT. Conservation calculation method was set to the Bayesian method. Evolutionary substitution model was selected as the Best Model (default value). Estimated conservation values were mapped onto the LRP1 dimer model and visualised as cartoon putty using PyMol Molecular Graphics System (Schroedinger, LLC).

## Acknowledgements

This work was supported by the Ministry of Science and Innovation of Spain, FPI grant PRE2021-100507 (G.M.T.); Grants I+D+I PID2020-119914RBI00 and PID2023-149206OB-I00 (G.B.), Grant PID2021-124297NB-C31 (G.F.) and PID2022-138109OB-I00 (L.R.P.) funded by MCIN/AEI/10.13039/501100011033 and European Union “ERDF A way to make Europe” (G.F.); the Alzheimer’s Association New to the field award (G.B.), European Research Council ChessTaG grant 769798 and MAIN (101123468) (G.B.); Agencia de Gestíon de Ayudas Universitaria y de Investigacíon-Beatriu de Pinos Fellowship (S.A.G.); Activitat científica dels grups de recerca de Catalunya (SGR-Cat 2021) (G.B.); Marco Basile has received the support of a fellowship from “la Caixa” Foundation (ID 100010434). The fellowship code is “LCF/BQ/DI22/11940010”. (M.B.); National Key R&D Program of China (2022YFC2009900) (X.T.). We thank the Spanish Network for Supercomputing (Red Española de Supercomputacion, RES) for the computational resources with the projects BCV-2023-3-0019.

## Supporting Information

**Figure S1.**
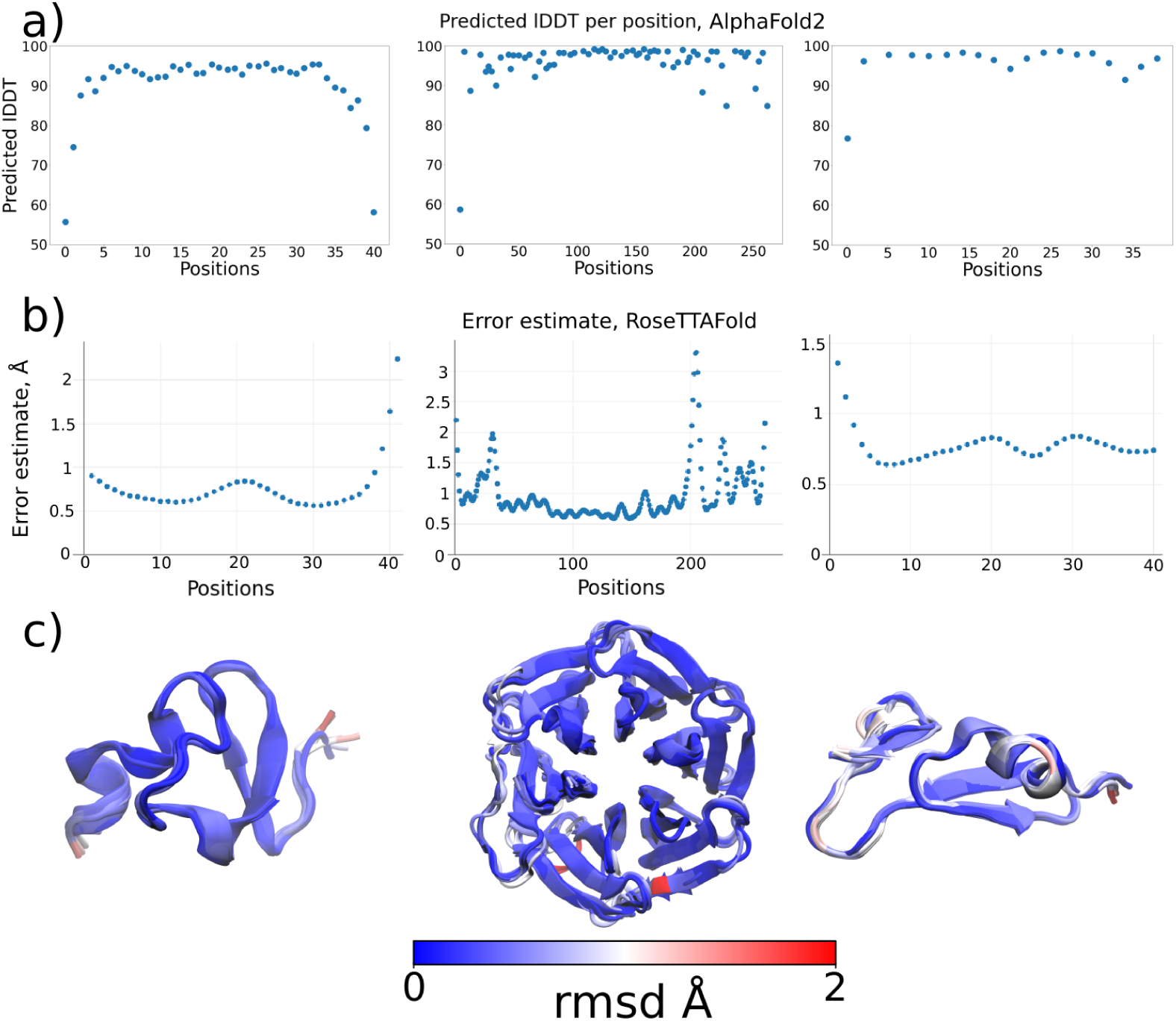
Evaluation of LRP1’s predicted motifs using structure predicting algorithms. a) Predicted local distance difference test lDDT per position of the AlphaFold2 prediction for each motif, from left to right: calcium-binding motif, β-propeller motif, EGF-like motif. Values above 90 correspond to very high confidence of the structure prediction. b) Error estimate from the RoseTTAFold prediction for each motif, following the order in (a). c) Structural alignment of the five RoseTTAFold and five AlphaFold2’s prediction of the three motifs, following the order in (a). The amino acids are highly super-imposed, demonstrating the two neural network’s prediction quality.

**Figure S2.**
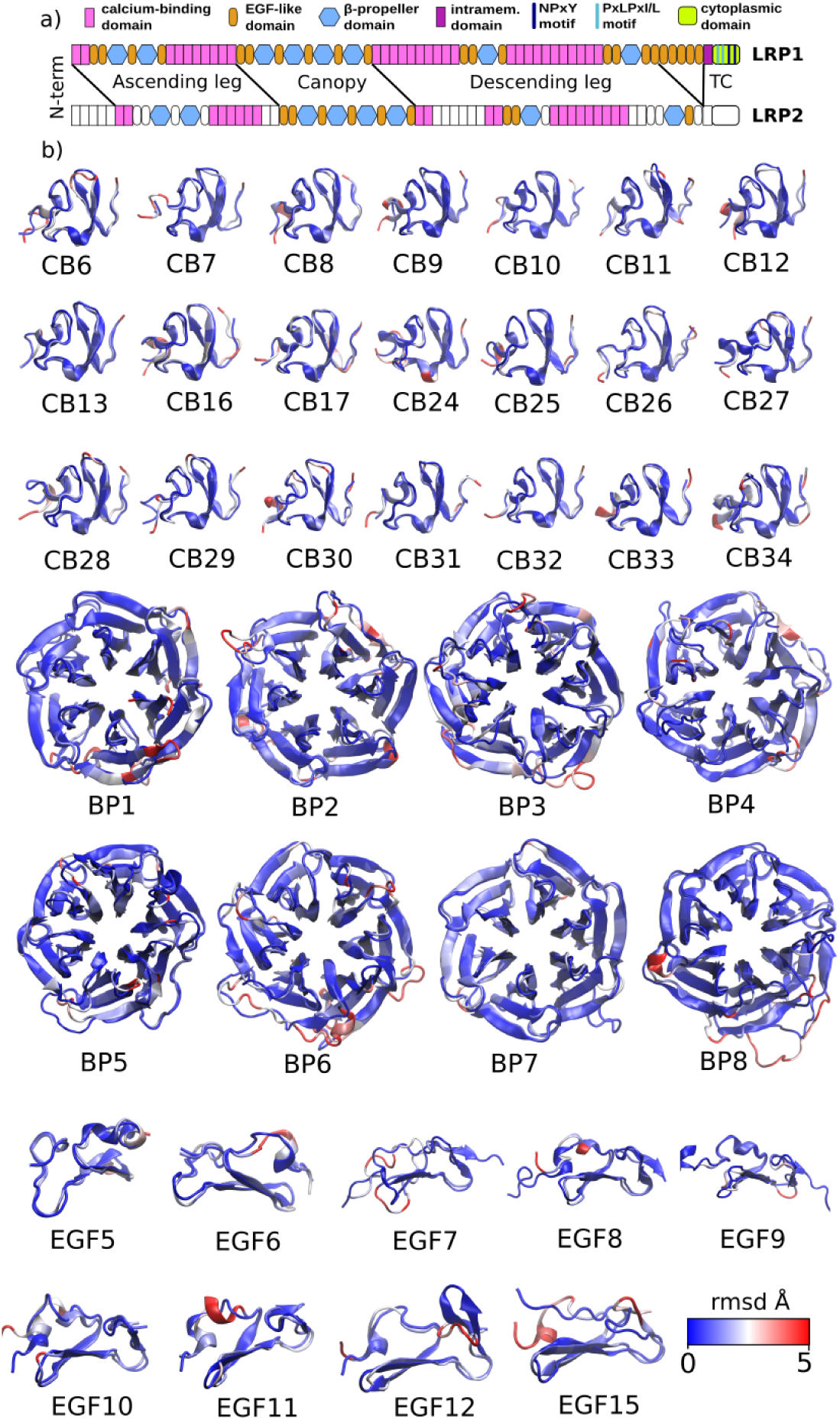
Evaluation of LRP1’s predicted motifs using structure predicting algorithms. a) Alignment of the LRP1 and LRP2 structural sequences. It is evident the similarity between the two motif sequences, if ignoring the first five CB units in LRP2 (counting from N-terminal) and the last five EGF-like motifs in LRP1. The motifs in white in LRP2 are the ones not used by 8EM4 for the analysis. b) Structural superimposition of the motifs from the cryo-TEM observation 8EM4 and the corresponding motif in the LRP1 neural network prediction (RoseTTAFold). Each couple satisfyingly superimpose, confirming the validity of the NN in the local prediction.

**Figure S3.**
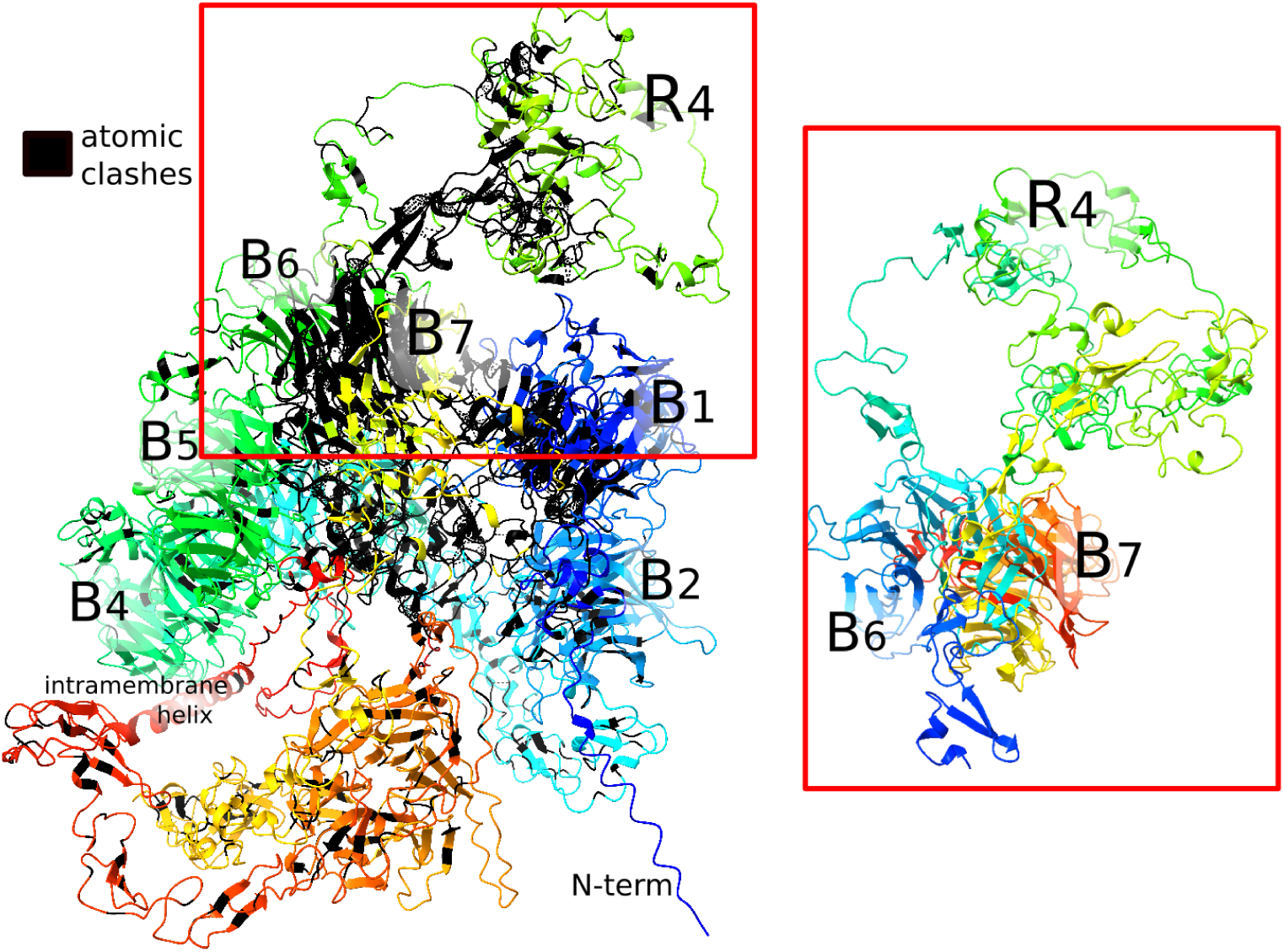
Evaluation of AlphaFold3 full LRP1 prediction. Prediction of the LRP1 full protein by AlphaFold3, shown as a cartoon and colored with rainbow colors from N terminus (blue) to C terminus (red). The N terminus and the intramembrane helix close to the C terminus are visible. LRP1 has a coiled conformation with many contacts (amino acids in black) and even a superimposition of the β-propellers 6 and 7, highlighted by the red square. The structure has not the required quality for practical uses.

**Figure S4.**
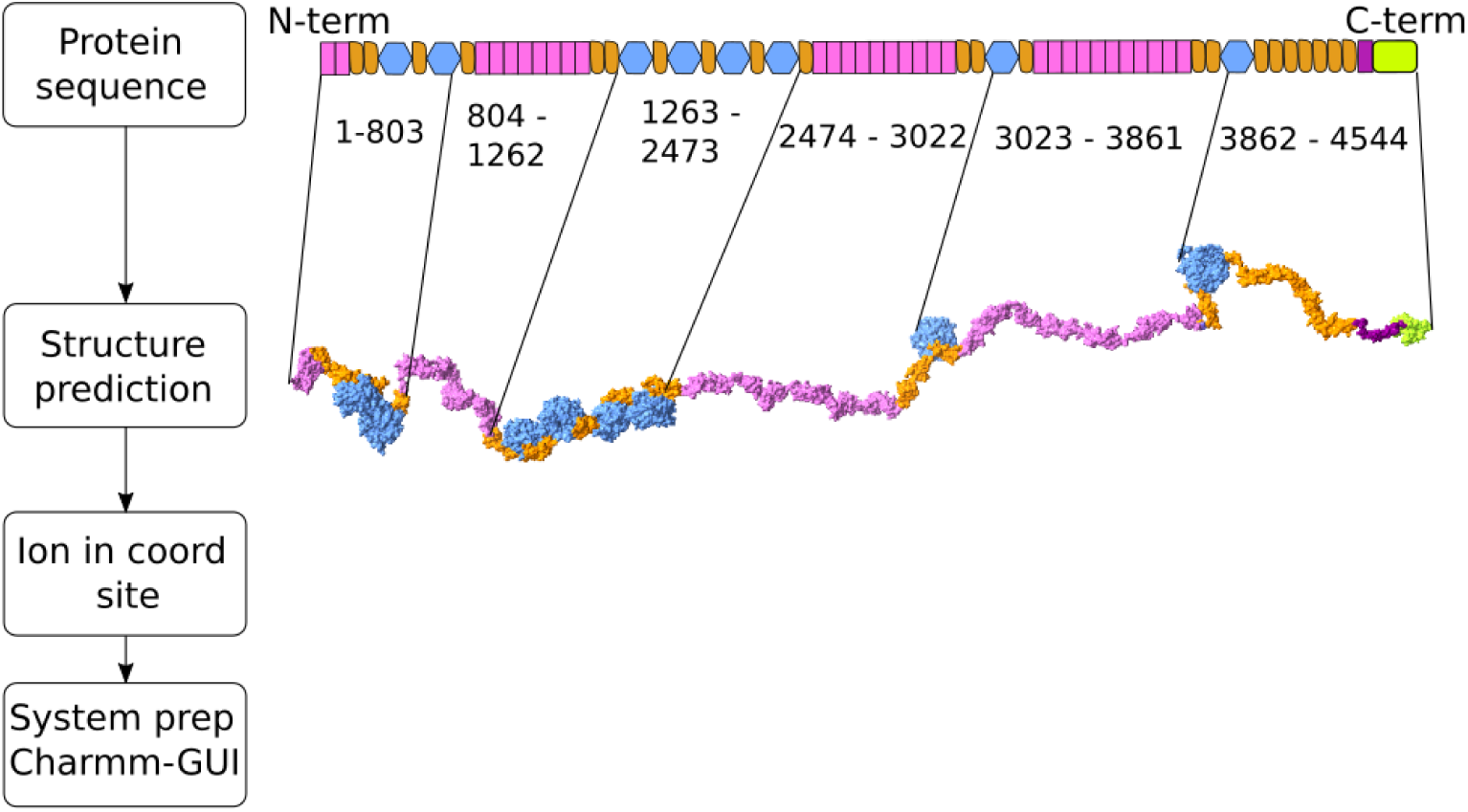
Production of a single LRP1 by joining of protein fragments. Protocol for the production of the LRP1 monomeric model. We first separated the protein sequence into six portions and produced for each a predicted structure using RoseTTAFold. The fragments were then joined and adjusted in the conformation shown here and in Fig. 3.b. We further added calcium ions to the coordination sites of the calcium-binding motifs and finally prepared the glycoprotein for molecular dynamics simulations using the Charmm-GUI web-server [32].

**Figure S5.**
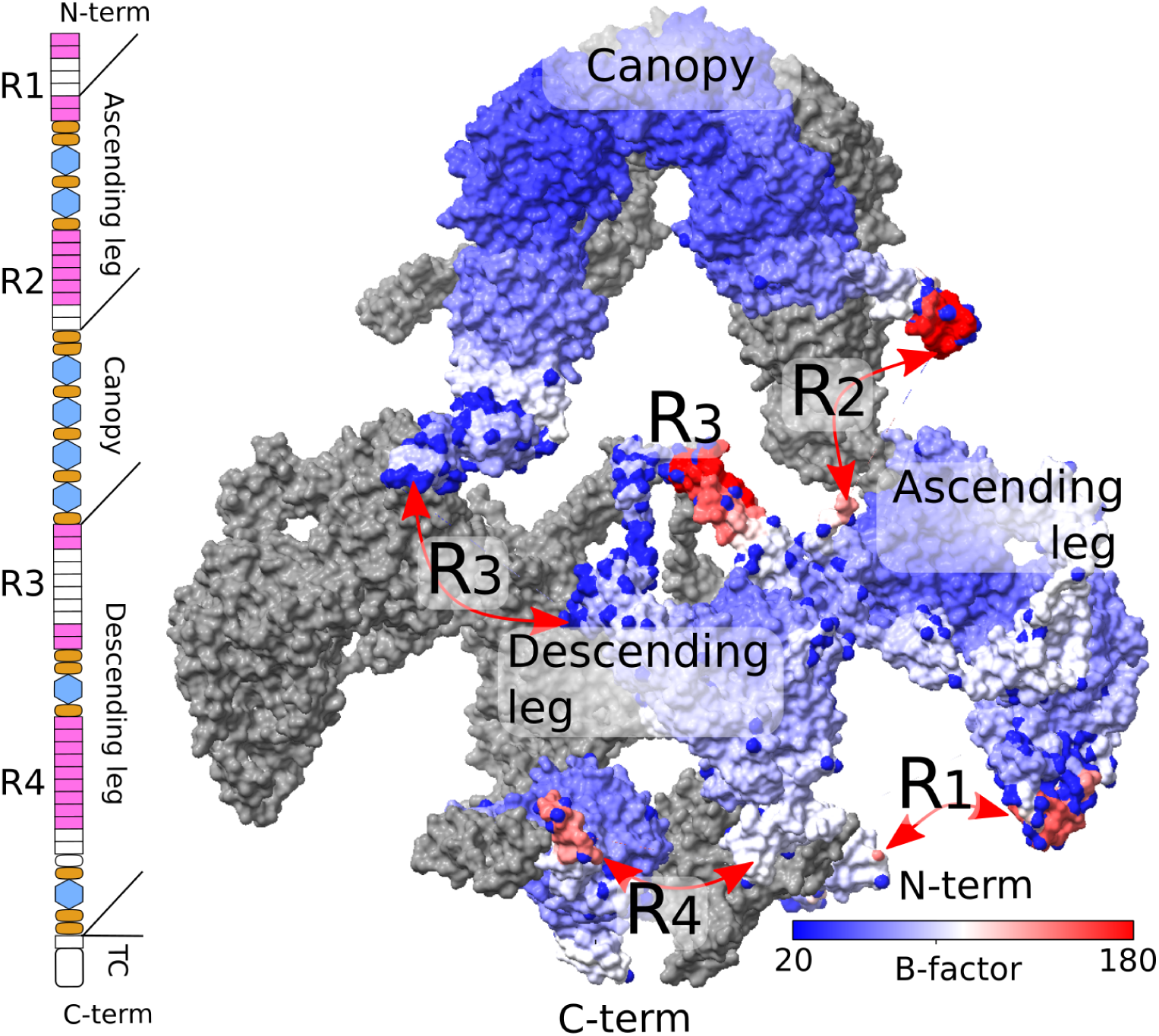
LRP2 cryo-TEM structure B-factor enhances the flexibility of specific domains. LRP2 at pH 7.5 (PDB 8EM4) obtained by the observations of Beenken et al [18]. The two proteins of the dimer are shown in vdW surface; the forefront monomer displays the atomic B-factor values, and the background is colored grey. The red indicates a greater variability of the atomic positions. Notably, the atoms in calcium-binding domains R1, R2, R3 and R4 are more prone to move; some portions of the domains are missing because the atomic positions were not defined with enough high resolution, highlighting an overall greater flexibility of the R domains compared with the rest of the protein. The white units of the scheme on the left show the missing units of 8EM4.

**Figure S6.**
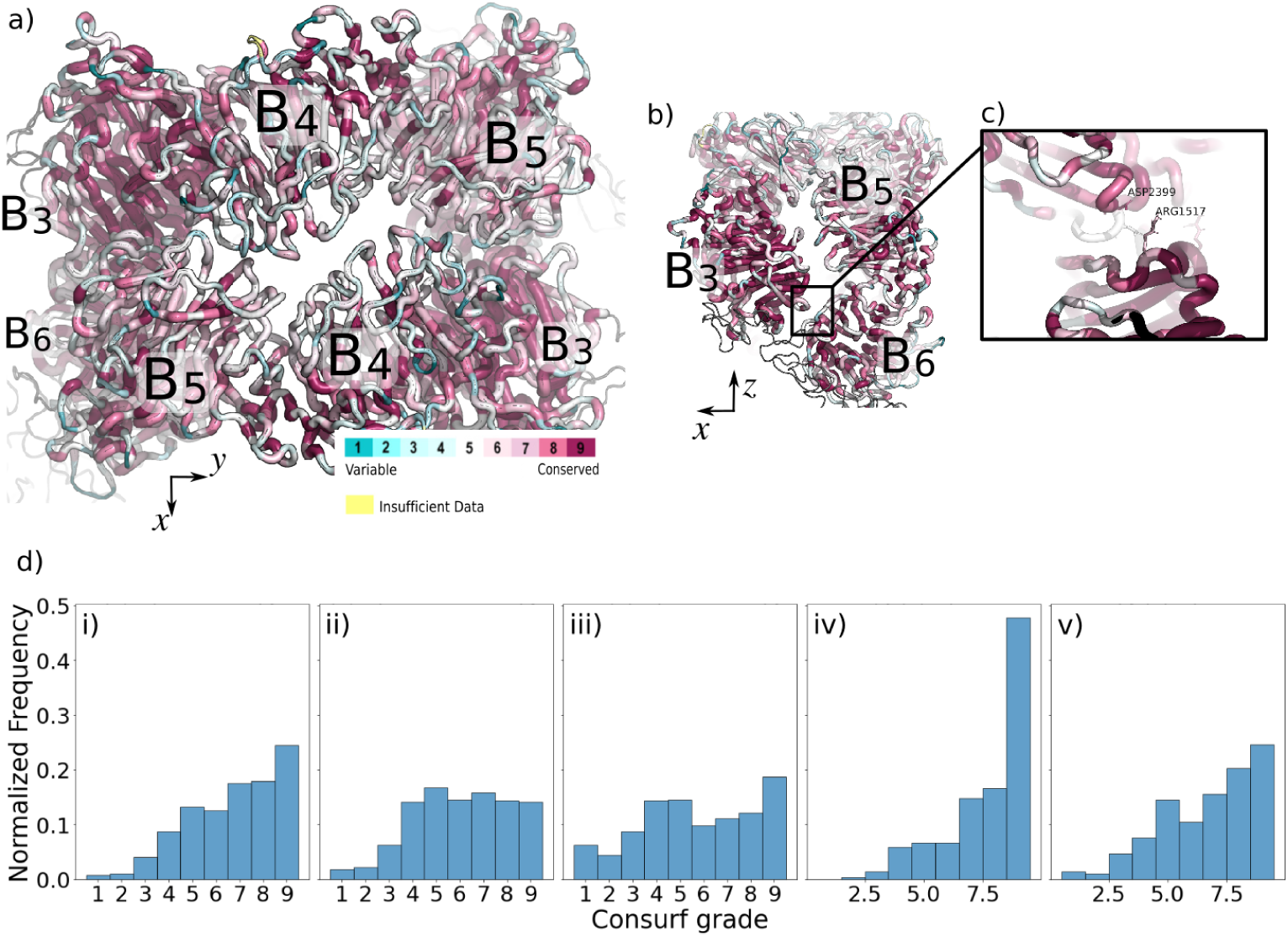
Evolutionary conservation. a) Top view of the LRP1 dimeric canopy. Amino acids are colored following the ConSurf [35] scheme (See Methods). High scores indicate the residue is highly conserved among identified homologue protein structures, indicating functional importance. A general trend is observed that core residues of the β-propellers are more conserved than the ones at the surface. Lower scores of surface residues indicate their non-functional role at the dimeric interface. b) The most conserved amino acids at the interface are between β-propellers 3 and 6. c) Detail of amino acids ASP2399 and ARG1517, the most conserved among the ones involved in hydrogen bonds (see Tab. 1). d) Histograms showing the probability density of finding a Consurf grade: i) in the canopy β-propellers (3 to 6); ii) in the β-propellers outside the canopy (1, 2, 7, 8); iii) in the full protein, excluding the canopy; iv) in β-propeller 3; v) in β-propeller 6.

## References

1. Steven L. Gonias and W. Marie Campana. Ldl receptor–related protein-1: A regulator of inflammation in atherosclerosis, cancer, and injury to the nervous system. The American Journal of Pathology, 184:18, 1 2014.

2. Nicola Potere, Marco Giuseppe Del Buono, Adolfo Gabriele Mauro, Antonio Abbate, and Stefano Toldo. Low density lipoprotein receptor-related protein-1 in cardiac inflammation and infarct healing. Frontiers in Cardiovascular Medicine, 6:51, 4 2019.

3. Hervé Emonard and Etienne Marbaix. Low-density lipoprotein receptor-related protein in metalloproteinase-mediated pathologies: recent insights. Metalloproteinases In Medicine, 2:9–18, 2 2015.

4. Dianaly T. Au, Dudley K. Strickland, and Selen C. Muratoglu. The ldl receptor-related protein 1: At the crossroads of lipoprotein metabolism and insulin signaling. Journal of Diabetes Research, 2017, 2017.

5. Mitsuru Shinohara, Masaya Tachibana, Takahisa Kanekiyo, and Guojun Bu. Thematic review series: Apoe and lipid homeostasis in alzheimer’s disease: Role of lrp1 in the pathogenesis of alzheimer’s disease: evidence from clinical and preclinical studies. Journal of Lipid Research, 58:1267, 2017.

6. Anna P. Lillis, Selen Catania Muratoglu, Dianaly T. Au, Mary Migliorini, Mi Jeong Lee, Susan K. Fried, Irina Mikhailenko, and Dudley K. Strickland. Ldl receptor-related protein-1 (lrp1) regulates cholesterol accumulation in macrophages. PLoS ONE, 10, 6 2015.

7. Chikako Nakajima, Philipp Haffner, Sebastian M. Goerke, Kai Zurhove, Giselind Adelmann, Michael Frotscher, Joachim Herz, Hans H. Bock, and Petra May. The lipoprotein receptor lrp1 modulates sphingosine-1-phosphate signaling and is essential for vascular development. Development, 141:4513–4525, 12 2014.

8. Qiang Liu, Justin Trotter, Juan Zhang, Melinda M. Peters, Hua Cheng, Jianxin Bao, Xianlin Han, Edwin J. Weeber, and Guojun Bu. Neuronal lrp1 knockout in adult mice leads to impaired brain lipid metabolism and progressive, age-dependent synapse loss and neurodegeneration. Journal of Neuroscience, 30:17068–17078, 12 2010.

9. Anna P. Lillis, Lauren B. Van Duyn, Joanne E. Murphy-Ullrich, and Dudley K. Strickland. The low density lipoprotein receptor-related protein 1: Unique tissue-specific functions revealed by selective gene knockout studies. Physiological reviews, 88:887, 7 2008.

10. T E Willnow, J M Moehring, N M Inocencio, T J Moehring, and J Herz. The low-density-lipoprotein receptor-related protein (lrp) is processed by furin in vivo and in vitro. The Biochemical Journal, 313 ( Pt 1):71–76, 1 1996.

11. Camilla De Nardis, Philip Lossl, Maartje Van Den Biggelaar, Pramod K. Madoori, Nadia Leloup, Koen Mertens, Albert J.R. Heck, Piet Gros, and Norma Allewell. Recombinant expression of the full-length ectodomain of ldl receptor-related protein 1 (lrp1) unravels phdependent conformational changes and the stoichiometry of binding with receptor-associated protein (rap). Journal of Biological Chemistry, 292:912–924, 1 2017.

12. M. Simonovic, K. Dolmer, W. Huang, D. K. Strickland, K. Volz, and P. G.W. Gettins. Calcium coordination and ph dependence of the calcium affinity of ligand-binding repeat cr7 from the lrp. comparison with related domains from the lrp and the ldl receptor. Biochemistry, 40:15127–15134, 12 2001.

13. Gabby Rudenko, Lisa Henry, Keith Henderson, Konstantin Ichtchenko, Michael S. Brown, Joseph L. Goldstein, and Johann Deisenhofer. Structure of the ldl receptor extracellular domain at endosomal ph. Science, 298:2353–2358, 12 2002.

14. Timothy A. Springer. An extracellular β-propeller module predicted in lipoprotein and scavenger receptors, tyrosine kinases, epidermal growth factor precursor, and extracellular matrix components. Journal of Molecular Biology, 283:837–862, 11 1998.

15. Hyesung Jeon, Wuyi Meng, Junichi Takagi, Michael J Eck, Timothy A Springer, and Stephen C Blacklow. Implications for familial hypercholesterolemia from the structure of the ldl receptor ywtd-egf domain pair. Nature Structural Biology, 8:499–504, 6 2001.

16. J. Herz, U. Hamann, S. Rogne, O. Myklebost, H. Gausepohl, and K. K. Stanley. Surface location and high affinity for calcium of a 500-kd liver membrane protein closely related to the ldl-receptor suggest a physiological role as lipoprotein receptor. The EMBO Journal, 7:4119–4127, 12 1988.

17. Daniela Passarella, Silvia Ciampi, Valentina Di Liberto, Mariachiara Zuccarini, Maurizio Ronci, Alessandro Medoro, Emanuele Foderà, Monica Frinchi, Donatella Mignogna, Claudio Russo, and Carola Porcile. Low-density lipoprotein receptor-related protein 8 at the crossroad between cancer and neurodegeneration. International Journal of Molecular Sciences 2022, Vol. 23, Page 8921, 23:8921, 8 2022.

18. Andrew Beenken, Gabriele Cerutti, Julia Brasch, Yicheng Guo, Zizhang Sheng, Hediye Erdjument-Bromage, Zainab Aziz, Shelief Y. Robbins-Juarez, Estefania Y. Chavez, Goran Ahlsen, Phinikoula S. Katsamba, Thomas A. Neubert, Anthony W.P. Fitzpatrick, Jonathan Barasch, and Lawrence Shapiro. Structures of lrp2 reveal a molecular machine for endocytosis. Cell, 186:821, 2 2023.

19. Carlos Spuch, Saida Ortolano, and Carmen Navarro. Lrp-1 and lrp-2 receptors function in the membrane neuron. trafficking mechanisms and proteolytic processing in alzheimer’s disease. Frontiers in Physiology, 3, 2012.

20. Sawako Goto, Akihisa Tsutsumi, Yongchan Lee, Michihiro Hosojima, Hideyuki Kabasawa, Koichi Komochi, Satoru Nagatoishi, Kazuya Takemoto, Kouhei Tsumoto, Tomohiro Nishizawa, Masahide Kikkawa, and Akihiko Saito. Cryo-em structures elucidate the multiligand receptor nature of megalin. Proceedings of the National Academy of Sciences of the United States of America, 121:e2318859121, 5 2024.

21. Ajit Varki. Biological roles of glycans. Glycobiology, 27:3–49, 1 2017.

22. Silvia Acosta-Gutiérrez, Joseph Buckley, Giuseppe Battaglia, S Acosta-Gutiérrez, J Buckley, and G Battaglia. The role of host cell glycans on virus infectivity: The sars-cov-2 case. Advanced Science, 10:2201853, 1 2023.

23. Atanu Acharya, Diane L. Lynch, Anna Pavlova, Yui Tik Pang, and James C. Gumbart. Ace2 glycans preferentially interact with sars-cov-2 over sars-cov. Chemical Communications, 57:5949–5952, 6 2021.

24. Kanika Arora, P. M. Sherilraj, K. A. Abutwaibe, Bharti Dhruw, and Shyam Lal Mudavath. Exploring glycans as vital biological macromolecules: A comprehensive review of advancements in biomedical frontiers. International Journal of Biological Macromolecules, 268:131511, 5 2024.

25. Junyang Chen, Xiang Pan, Aroa Duro-Castano, Huawei Cai, Bin Guo, Xiqin Liu, Yifan Yu, Su Lui, Kui Luo, Bowen Ke, Lorena Ruiz Perez, Xiawei Wei, Qiyong Gong, Xiaohe Tian, and Giuseppe Battaglia. Multivalent targeting of blood-brain barrier lrp1 for neurovascular recovery therapy for alzheimer’s disease. bioRxiv, page 2024.05.06.592767, 5 2024.

26. Xiaohe Tian, Diana M. Leite, Edoardo Scarpa, Sophie Nyberg, Gavin Fullstone, Joe Forth, Diana Matias, Azzurra Apriceno, Alessandro Poma, Aroa Duro-Castano, Manish Vuyyuru, Lena Harker-Kirschneck, Andela S^̌^aríc, Zhongping Zhang, Pan Xiang, Bin Fang, Yupeng Tian, Lei Luo, Loris Rizzello, and Giuseppe Battaglia. On the shuttling across the blood-brain barrier via tubule formation: Mechanism and cargo avidity bias. Science Advances, 6:4397–4424, 11 2020.

27. John Jumper, Richard Evans, Alexander Pritzel, Tim Green, Michael Figurnov, Olaf Ronneberger, Kathryn Tunyasuvunakool, Russ Bates, Augustin Žídek, Anna Potapenko, Alex Bridgland, Clemens Meyer, Simon A.A. Kohl, Andrew J. Ballard, Andrew Cowie, Bernardino Romera-Paredes, Stanislav Nikolov, Rishub Jain, Jonas Adler, Trevor Back, Stig Petersen, David Reiman, Ellen Clancy, Michal Zielinski, Martin Steinegger, Michalina Pacholska, Tamas Berghammer, Sebastian Bodenstein, David Silver, Oriol Vinyals, Andrew W. Senior, Koray Kavukcuoglu, Pushmeet Kohli, and Demis Hassabis. Highly accurate protein structure prediction with alphafold. Nature 2021 596:7873, 596:583–589, 7 2021.

28. Minkyung Baek, Frank DiMaio, Ivan Anishchenko, Justas Dauparas, Sergey Ovchinnikov, Gyu Rie Lee, Jue Wang, Qian Cong, Lisa N. Kinch, R. Dustin Schaeffer, Claudia Millán, Hahnbeom Park, Carson Adams, Caleb R. Glassman, Andy DeGiovanni, Jose H. Pereira, Andria V. Rodrigues, Alberdina A. Van Dijk, Ana C. Ebrecht, Diederik J. Opperman, Theo Sagmeister, Christoph Buhlheller, Tea Pavkov-Keller, Manoj K. Rathinaswamy, Udit Dalwadi, Calvin K. Yip, John E. Burke, K. Christopher Garcia, Nick V. Grishin, Paul D. Adams, Randy J. Read, and David Baker. Accurate prediction of protein structures and interactions using a three-track neural network. Science, 373:871–876, 8 2021.

29. Shankar Kumar, John M. Rosenberg, Djamal Bouzida, Robert H. Swendsen, and Peter A. Kollman. The weighted histogram analysis method for free-energy calculations on biomolecules. i. the method. Journal of Computational Chemistry, 13:1011–1021, 10 1992.

30. Fábio Madeira, Matt Pearce, Adrian R.N. Tivey, Prasad Basutkar, Joon Lee, Ossama Edbali, Nandana Madhusoodanan, Anton Kolesnikov, and Rodrigo Lopez. Search and sequence analysis tools services from embl-ebi in 2022. Nucleic Acids Research, 50:W276–W279, 7 2022.

31. Benjamin Webb and Andrej Sali. Comparative protein structure modeling using modeller. Current Protocols in Bioinformatics, 54:5.6.1–5.6.37, 6 2016.

32. Sunhwan Jo, Taehoon Kim, Vidyashankara G. Iyer, and Wonpil Im. Charmm-gui: A web-based graphical user interface for charmm. Journal of Computational Chemistry, 29:1859–1865, 8 2008.

33. William S. York, Raja Mazumder, Rene Ranzinger, Nathan Edwards, Robel Kahsay, Kiyoko F. Aoki-Kinoshita, Matthew P. Campbell, Richard D. Cummings, Ten Feizi, Maria Martin, Darren A. Natale, Nicolle H. Packer, Robert J. Woods, Gaurav Agarwal, Sena Arpinar, Sanath Bhat, Judith Blake, Leyla Jael Garcia Castro, Brian Fochtman, Jeffrey Gildersleeve, Radoslav Goldman, Xavier Holmes, Vinamra Jain, Sujeet Kulkarni, Rupali Mahadik, Akul Mehta, Reza Mousavi, Sandeep Nakarakommula, Rahi Navelkar, Nagarajan Pattabiraman, Michael J. Pierce, Karen Ross, Preethi Vasudev, Jeet Vora, Tatiana Williamson, and Wenjin Zhang. Glygen: Computational and informatics resources for glycoscience. Glycobiology, 30:72–73, 2 2020.

34. Thomas Lemmin and Cinque Soto. Glycosylator: A python framework for the rapid modeling of glycans. BMC Bioinformatics, 20:1–7, 10 2019.

35. Fabian Glaser, Tal Pupko, Inbal Paz, Rachel E. Bell, Dalit Bechor-Shental, Eric Martz, and Nir Ben-Tal. Consurf: identification of functional regions in proteins by surface-mapping of phylogenetic information. *Bioinformatics (Oxford*, England), 19:163–164, 1 2003.

36. Kathryn A. Quinn, Philip G. Grimsley, Yang Ping Dai, Michael Tapner, Colin N. Chesterman, and Dwain A. Owensby. Soluble low density lipoprotein receptor-related protein (lrp) circulates in human plasma. Journal of Biological Chemistry, 272:23946–23951, 9 1997.

37. Kathryn A. Quinn, Victoria J. Pye, Yang Ping Dai, Colin N. Chesterman, and Dwain A. Owensby. Characterization of the soluble form of the low density lipoprotein receptor-related protein (lrp). Experimental Cell Research, 251:433– 441, 9 1999.

38. Eric F. Pettersen, Thomas D. Goddard, Conrad C. Huang, Gregory S. Couch, Daniel M. Greenblatt, Elaine C. Meng, and Thomas E. Ferrin. Ucsf chimera—a visualization system for exploratory research and analysis. Journal of Computational Chemistry, 25:1605–1612, 10 e.

39. Mengyuan He, Xiangxiang Zhou, and Xin Wang. Glycosylation: mechanisms, biological functions and clinical implications. Signal Transduction and Targeted Therapy 2024 9:1, 9:1–33, 8 2024.

40. Yingwei Hu, Punit Shah, David J. Clark, Minghui Ao, and Hui Zhang. Reanalysis of global proteomic and phosphoproteomic data identified a large number of glycopeptides. Analytical Chemistry, 90:8065–8071, 7 2018.

41. T. Mamie Lih, Kyung Cho Cho, Michael Schnaubelt, Yingwei Hu, and Hui Zhang. Integrated glycoproteomic characterization of clear cell renal cell carcinoma. Cell Reports, 42, 5 2023.

42. Peter Eastman, Jason Swails, John D. Chodera, Robert T. McGibbon, Yutong Zhao, Kyle A. Beauchamp, Lee Ping Wang, Andrew C. Simmonett, Matthew P. Harrigan, Chaya D. Stern, Rafal P. Wiewiora, Bernard R. Brooks, and Vijay S. Pande. Openmm 7: Rapid development of high performance algorithms for molecular dynamics. PLOS Computational Biology, 13:e1005659, 7 2017.

43. Zhijun Zhang, Xinzijian Liu, Kangyu Yan, Mark E. Tuckerman, and Jian Liu. Unified efficient thermostat scheme for the canonical ensemble with holonomic or isokinetic constraints via molecular dynamics. Journal of Physical Chemistry A, 123:6056–6079, 5 2019.

44. A. D. MacKerell, D. Bashford, M. Bellott, R. L. Dunbrack, J. D. Evanseck, M. J. Field, S. Fischer, J. Gao, H. Guo, S. Ha, D. Joseph-McCarthy, L. Kuchnir, K. Kuczera, F. T.K. Lau, C. Mattos, S. Michnick, T. Ngo, D. T. Nguyen, B. Prodhom, W. E. Reiher, B. Roux, M. Schlenkrich, J. C. Smith, R. Stote, J. Straub, M. Watanabe, J. Wiórkiewicz-Kuczera, D. Yin, and M. Karplus. All-atom empirical potential for molecular modeling and dynamics studies of proteins. Journal of Physical Chemistry B, 102:3586–3616, 4 1998.

45. Hai Nguyen, Daniel R. Roe, and Carlos Simmerling. Improved generalized born solvent model parameters for protein simulations. Journal of chemical theory and computation, 9:2020–2034, 4 2013.

46. Mark James Abraham, Teemu Murtola, Roland Schulz, Szilárd Páll, Jeremy C. Smith, Berk Hess, and Erik Lindah. Gromacs: High performance molecular simulations through multi-level parallelism from laptops to supercomputers. SoftwareX, 1-2:19–25, 9 2015.

47. Jing Huang, Sarah Rauscher, Grzegorz Nawrocki, Ting Ran, Michael Feig, Bert L. De Groot, Helmut Grubmüller, and Alexander D. MacKerell. Charmm36m: An improved force field for folded and intrinsically disordered proteins. Nature methods, 14:71, 12 2017.

48. Jochen S. Hub, Bert L. De Groot, and David Van Der Spoel. G-whams-a free weighted histogram analysis implementation including robust error and autocorrelation estimates. Journal of Chemical Theory and Computation, 6:3713–3720, 12 2010.

49. Naveen Michaud-Agrawal, Elizabeth J. Denning, Thomas B. Woolf, and Oliver Beckstein. Mdanalysis: A toolkit for the analysis of molecular dynamics simulations. Journal of Computational Chemistry, 32:2319–2327, 7 2011.

50. William Humphrey, Andrew Dalke, and Klaus Schulten. Vmd: Visual molecular dynamics. Journal of Molecular Graphics, 14:33–38, 2 1996.

51. Aihua Zhang, Hua Yu, Chunhong Liu, and Chen Song. The ca2+ permeation mechanism of the ryanodine receptor revealed by a multi-site ion model. Nature Communications 2020 11:1, 11:1–10, 2 2020.

